# Focus-ISM for Sharp and Gentle Super-Resolved Microscopy

**DOI:** 10.1101/2022.04.28.489892

**Authors:** Giorgio Tortarolo, Alessandro Zunino, Francesco Fersini, Marco Castello, Simonluca Piazza, Colin J.R. Sheppard, Paolo Bianchini, Alberto Diaspro, Sami Koho, Giuseppe Vicidomini

**Affiliations:** Molecular Microscopy and Spectroscopy, Istituto Italiano di Tecnologia, Genoa, Italy; DIBRIS, University of Genoa, Genoa, Italy; Nanoscopy, Istituto Italiano di Tecnologia, Genoa, Italy; DIFI, University of Genoa, Genoa, Italy

**Author notes:** These authors contributed equally to this work (Giorgio Tortarolo, Alessandro Zunino). Laboratory of Experimental Biophysics, EPFL, Lausanne, Switzerland.

## Abstract

Super-resolution microscopy is routinely used for fixed and thin samples, while its feasibility for imaging live and thick samples is still limited. In the case of stimulated emission depletion (STED) microscopy, the high-intensity illumination required to achieve effective sub-diffraction resolution can introduce photo-damage, thus reducing the compatibility of the technique with live-cell imaging. Moreover, the out-of-focus fluorescence background may overcome the often faint signal stemming from the focal point, thus constraining imaging to thin samples. Here, we combined STED microscopy with image-scanning microscopy (ISM) to mitigate these limitations without any practical disadvantages. We first enhanced a laser scanning microscope (LSM) by introducing a detector array, hence providing access to a set of additional spatial information that is not available with a typical single-element detector. Then, we exploited this extended dataset to implement focus-ISM, a novel method that relaxes the high-intensity requirement of STED microscopy and removes the out-of-focus background. Additionally, we generalized the focus-ISM method to conventional LSM, namely without a STED beam. The proposed approach requires minimal architectural changes compared with conventional STED microscopes but provides substantial advantages for live and thick sample imaging while maintaining all compatibility with all recent advances in STED and confocal microscopy. As such, focus-ISM represents an essential step towards a universal super-resolved LSM technique for subcellular imaging.

## Introduction

The vast family of fluorescence optical microscopy techniques stands as an invaluable tool for addressing various biological questions, enabling the dynamical observation of bio-molecular processes in living cells – on their own or as part of a whole organism. Within this family, super-resolution (SR) microscopy techniques have opened up new exciting perspectives by overcoming the *classical* diffraction limit of spatial resolution – about half the wavelength of the fluorescence light [1, 2]. Confocal laser scanning-microscopy (CLSM) [3] was one of the first SR methods – at least theoretically. By illuminating the sample point-by-point and collecting the fluorescence light through a small pinhole, the spatial resolution could be improved by up to a factor of two below the *classical* diffraction limit – as for structured illumination microscopy (SIM). However, a closed pinhole leads to a strong reduction in signal-to-noise ratio (SNR), effectively precluding the resolution enhancement in a real-life scenario. Indeed, the historical success of CLSM is mostly due to its optical sectioning capability, rather than for the resolution improvements it theoretically provides. In more recent times, starting from the early 2000s, new SR microscopy concepts (referred to as diffraction-unlimited microscopy or nanoscopy) have not only surpassed the classical resolution limit, but also enabled theoretically unlimited resolution [4]. Stimulated emission depletion (STED) microscopy [5, 6] is one of the most versatile of such nanoscopy techniques, since it can attain multi-species and fast imaging with tunable resolution – from the *classical* diffraction-limit down to a theoretical molecular resolution. A STED microscope shares similar laser-scanning architecture to CLSM, but the Gaussian excitation beam is co-aligned with a second, vortex, beam – known as the STED beam, and both beams are scanned across the sample. Because the STED beam induces stimulated emission and depletes the fluorescence signal from the peripheral region of the excitation spot, the effective fluorescent region reduces in size well below the diffraction limit. Notably, the higher the STED beam intensity, the smaller the effective fluorescent region, and ultimately the better the spatial resolution. However, high STED beam intensity may cause photo-bleaching and ultimately photo-toxicity to the samples. Additionally, thick samples may emit out-of-focus fluorescence background decreasing the contrast of faint infocus fluorescence signals. The contrast further reduces if the STED beam excites directly the specimen, thus generating additional anti-Stokes fluorescence background. In other words, increasing the optical resolution comes at the cost of photo-damaging the sample and reducing the signal-to-background ratio (SBR), hence hindering the feasibility for long-term STED imaging in living cells and STED imaging in thick samples, respectively. Several STED microscopy implementations have been proposed to mitigate the photo-damage and the SBR reduction [7]: Time-resolved [8, 9, 10, 11, 12, 13] and subtraction methods [14, 15] remove the incomplete depletion background, aiming to reduce the STED beam intensity necessary to achieve a target resolution. Tomographic STED microscopy [16] obtains comparable intensity reductions by fusing multiple STED images collected with efficient two-dimensional STED beam intensity distributions. Synchronous detection schemes [10, 17, 11, 18] remove the anti-Stokes background. Adaptive smart scanning schemes [19, 20, 21] decrease the overall specimen illumination. Tailored three-dimensional STED beam intensity distributions [22, 23] chop the out-of-focus background. Among these discussed techniques, only time-resolved STED microscopy has now become a gold standard, being implemented in all commercial systems. The reasons behind this success lies in the simplicity of the technique: it requires only to register the temporal dynamics of the fluorescent signal, adding an additional temporal dimension to the image dataset. The other approaches provide great benefits but require a significant increase in technical complexity, leaving the scientific community still in need of easy strategies to improve the compatibility of STED microscopy with live-cell and thick samples.

More recently, image scanning microscopy (ISM) has effectively transformed CLSM into a SR technique. Such revolution has been made possible by substituting the typical single-element detector with a detector array, for which each element acts as an individual pinhole. In this case, no fluorescence signal is lost – since each detector element contributes to the collected signal – and the small size of the individual detector elements guarantees the resolution enhancement. In detail, the detector array records a bi-dimensional micro-image *i*(***x**_d_*|***x**_s_*), with ***x**_d_* = (*x_d_, y_d_*) the detector space, for each point of the scanning space ***x**_s_* = (*x_s_,y_s_*), effectively adding two additional spatial dimensions to the conventional image dataset. This feature justifies the name of the method as ISM. The two dimensions are related to a sample space by ***x*** = ***x**_d_* – ***x**_s_*. This additional information enable reconstructing via pixel-reassignment (PR) a sample’s image with a resolution enhancement close to the two-fold theoretical limit of of CLSM, without sacrificing SNR. Theoretically introduced by Sheppard in the 80s [24], and experimentally demonstrated by Enderlein in 2010 [25] with a conventional camera (~ kHz frame-rate), ISM became mainstream only with the introduction of fast (~ MHz frame-rate) detector arrays, such as the AiryScan [26] and single-photon avalanche diode (SPAD) array detectors [27, 28]: these effective super-resolved ISM implementations showed compatibility with multispecies, three-dimensional, and two-photon excitation imaging. At the same time, the SNR enhancement allows reduction of the excitation beam intensity, improving compatibility with live-cell imaging. Furthermore, detector arrays allow the combination of ISM with fluorescence lifetime imaging to increase further the information content, and with fluorescence photon-coincidences [29] or fluorescence fluctuations [30] imaging to boost the resolution beyond the twofold factor, and more generally to open up the possibility of diffraction-unlimited ISM. However, so far, these approaches require a long pixel-dwell time (≥ ms) to achieve high SNR, while obtaining small resolution enhancement, thus limiting their practical applications. Another class of approaches which may overcome the two-fold resolution enhancement of PR starting from the same ISM dataset is image deconvolution. Notably, in the 80s, Bertero [31] proposed a deconvolution method which works on the micro-images to achieve twice the resolution power of a conventional laser-scanning microscope.

In this work we show that ISM can be combined with STED microscopy to achieve a universal and versatile diffraction-unlimited SR technique. In particular, we introduce a STED-ISM implementation based on an asynchronous-readout SPAD array detector (Fig. 1a). Our architecture works at reduced STED beam intensity and rejects out-of-focus background, while maintaining compatibility with all the STED implementations mentioned above, and in particular with time-resolved STED microscopy. The detector array enables the acquisition for each scan point of a small wide-field image – named micro-image – of the excited region (Fig. 1b). The micro-images in general contain information on the lateral and axial structure of the sample, enabling the reconstruction of an image with enhanced content. In detail, (i) we implemented the adaptive pixel-reassignment (APR) method (Fig. 1c) to improve the spatial resolution while keeping a relatively low STED beam intensity, thus potentially reducing photo-damage; (ii) we introduced a classification method to separate the in-focus from the out-of-focus background signals, thus improving optical-sectioning.

**Figure 1:**
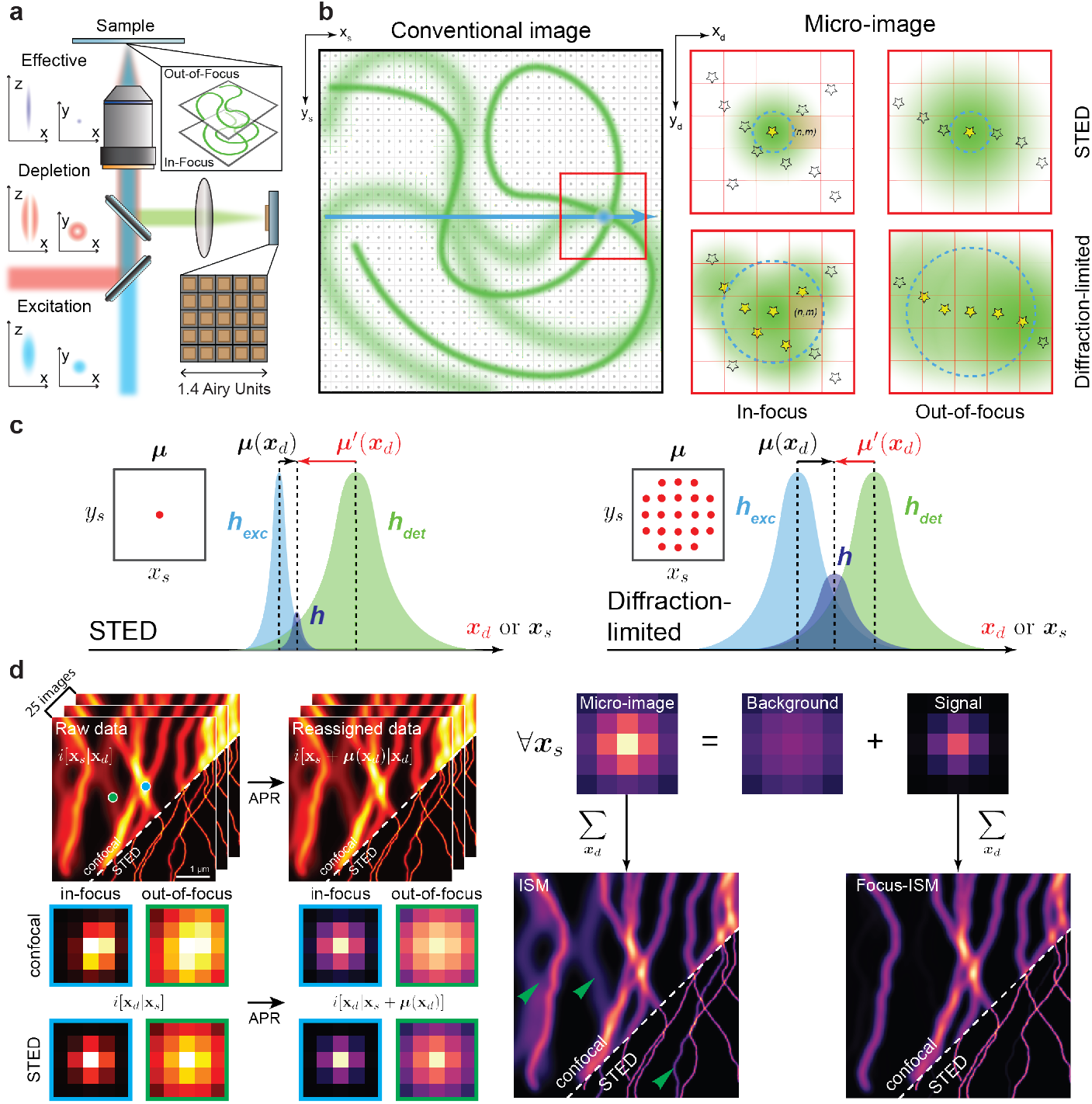
Focus-ISM imaging. In **a**, we present a sketch of the STED-ISM microscope, equipped with a 5 × 5 detector array. In **b**, we present a sketch of the image formation process in our setup. The sample – represented with an *in-focus* (sharp) and an *out-of-focus* (blurred) component – is raster scanned with the excitation beam. Each scan point (dark square) is related to a micro-image of the illuminated region (red square). The blue circle and the green halo represent, respectively, the excitation PSF and the emission of the excited fluorophores (yellow stars). In **c**, we depict the (adaptive) pixel-reassignment concept for a specific detector element (*n,m*). The shift-vector ***μ*** of a scanned image is the maximum position of the product of excitation and detection PSF. The higher is the STED power, the smaller is the excitation PSF, the shorter is the shift-vector. We also show the micro-image shift-vector ***μ***′. In **d**, we present the steps of focus-ISM reconstruction. The APR algorithm registers the scanned-images of the ISM dataset. APR affects also the micro-images, encoding uniquely the axial information of the sample. The Focus-ISM algorithm classifies the photons of each post-APR micro-image either as background or signal. Summing the pixels of the signal micro-images generates the Focus-ISM image. The green arrows indicate the out-of-focus filaments, no longer present in the focus-ISM image.

More generally, we investigated in detail the effects of the APR operation on different views of the multidimensional ISM data-set: whereas the post-APR scanned images are key for the ISM reconstruction, we demonstrate here that the post-APR micro-images directly encode the sample’s axial position for any STED intensity. We exploited this critical insight to develop focus-ISM, a novel two-step algorithm which first implements the well-know APR method and then a classification algorithm on the post-APR micro-images to extract the in-focus signal (Fig. 1d).

We tested the new focus-ISM method on calibration samples and fixed/living cells, achieving sharper STED microscopy images at any STED beam intensity. While the benefits of STED-ISM are reduced in the case of a high STED beam intensity, the focus-STED-ISM method improves optical-sectioning at any STED beam intensity – also in the limiting case of the simple confocal modality. Lastly, we further improved the contrast and the quality of the reconstructed images by means of a multi-image deconvolution algorithm, incorporating the out-of-focus background obtained thanks to focus-ISM as prior information.

## Results

### Reducing Intensity with Adaptive Pixel-Reassignment ISM

In this Section, we demonstrate how the concept of pixel reassignment can be efficiently combined with STED microscopy to achieve a target resolution at lower STED beam intensity. Our setup is a custom STED microscopy setup incorporating an asynchronous read-out SPAD array detector (Suppl. Fig. S1). The detector array enables the acquisition of a multi-dimensional dataset that can be regarded both as a collection of micro-images *i*(***x**_d_*|***x**_s_*) – one for each scan point ***x**_s_* – or as a collection of scanned-images *i*(***x**_s_*|***x**_d_*) – one for each detector element at position ***x**_d_*. The former perspective is the one adopted to apply the PR approach in the opto-mechanical ISM implementations [32] and, more generally, in the original PR idea (Supplementary Note 1). In ISM, the photons collected by each pixel of the detector are shifted from their most likely origin ***x*** = ***x**_s_* – ***x**_d_* + ***μ***′(***x**_d_*). Assuming no Stokes-shift and Gaussian point spread functions (PSFs), namely the distribution of light on the sample (excitation PSF) and on the detector (detection PSF), the so-called micro-image shift-vector ***μ***′(***x**_d_*) equals ***x**_d_*/2 and PR can be implementing by demagnifying twice the micro-image. More recently, we introduced the concept of APR to generalise PR to any imaging conditions [33, 34]. The APR approach regards the ISM dataset from the scanned-images point of view *i*(***x**_s_*|***x**_d_*) which is key for understanding the STED-ISM image formation. The images *i*(***x**_s_*|***x**_d_*) are mutually shifted by the quantities known as the scanned-image shift-vectors ***μ***(***x**_d_*). In short, the APR method registers the scanned-images and generates the ISM result, with enhanced resolution and SNR. The APR algorithm calculates the shift-vectors from the data by finding the shift that maximizes the similarity with a reference image, in our case the one generated by the central element of the detector array. For this reason, the APR method adaptively finds the best estimates of the shift-vectors directly from the images, taking inherently into account non-idealities – such as the-Stokes shift or optical aberrations – without any theoretical assumption. This aspect is of paramount importance for STED-ISM, because the value of the shift-vectors strongly depends on the STED power, making digital ISM the only technique compatible with STED microscopy.

Here, we show the benefits of applying the APR method directly to the raw STED-ISM dataset. For increasing STED beam intensities, we calculated the in-focus PSFs of all the scanned-images (Fig. 2a), the relative shift-vectors (Fig. 2b), and the in-focus PSFs of the conventional STED and of the STED-ISM image, reconstructed with the APR method (Fig. 2c). We obtained the conventional STED images by summing all the scanned-images, thus discarding the micro-image information, as it would happen with a single-element detector. The shift-vectors strongly depend on the STED beam intensity: the higher the depletion power (equivalently, the saturation factor), the smaller the effective fluorescent spot, and the shorter the shift-vectors. Based on the APR concept, the shift-vectors reflect the maximum position of the PSFs of each scanned image. Since the excitation PSF shrinks down to a single point for increasing STED beam intensity, its product with any detection PSF shrinks as well and its maximum position approaches the optical axis. In other words, in the case of high STED beam intensities, all scanned PSFs depend mainly on the effective excitation PSF and the influence of the detection PSF becomes negligible. Thus, the scanned-images vary in SNR but no longer in position. This insight shows that the adaptive pixel reassignment operation is beneficial for STED microscopy mainly for a specific range of STED intensities. More precisely, the spatial resolution and SNR of the STED-ISM reconstruction are improved with respect to the conventional STED counterparts for mild STED beam intensities (Fig. 2c, Suppl. Fig. S2). High STED beam intensities lead to very short shift-vectors and ultimately to negligible benefits from the APR method.

**Figure 2:**
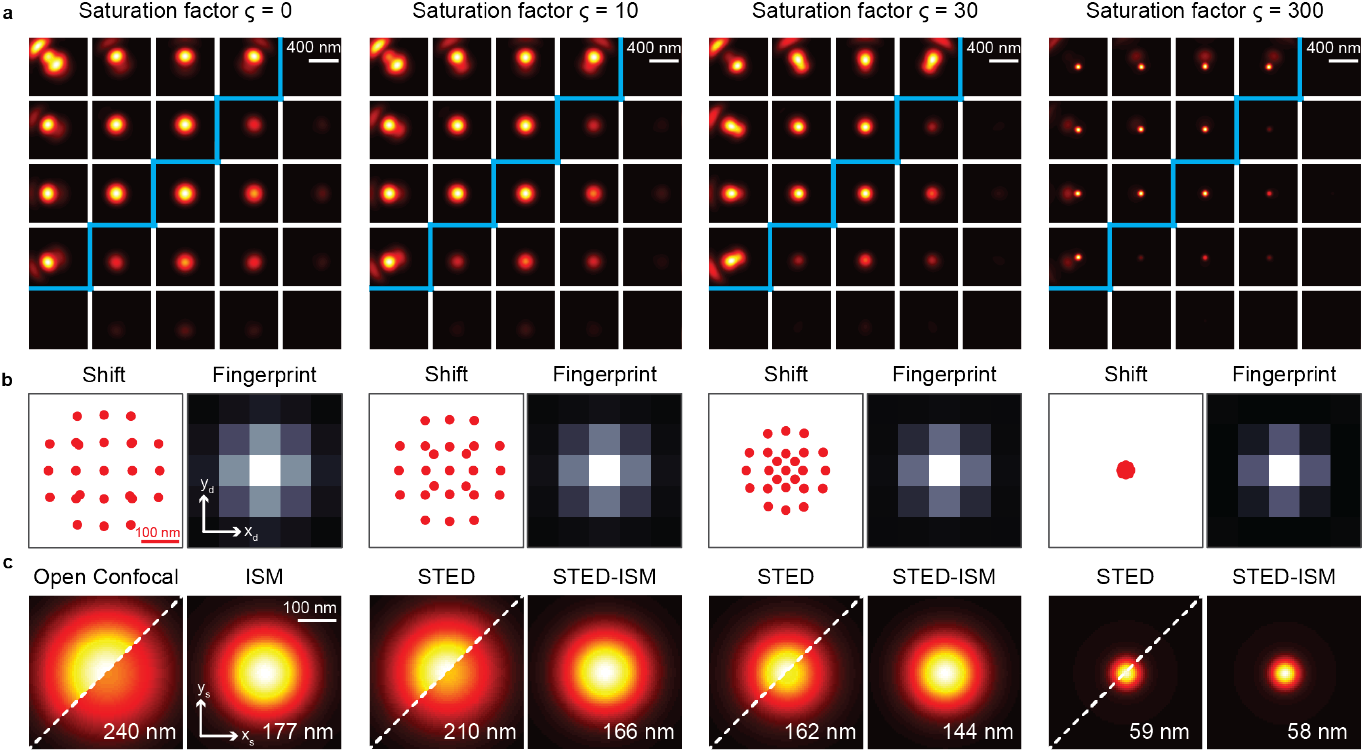
STED-ISM principle. In **a**, we show simulated images of the PSF *h*(***x**_s_*|***x**_d_*) for each detector element. In **b**, we show the *shift-vectors*, calculated using the adaptive pixel reassignment algorithm, and the *fingerprint*, calculated by summing all the ***x**_s_* points of the above images. In **c**, we compare the PSF obtained with a single-element detector (open pinhole configuration) with the PSF reconstructed with the ISM method. The upper diagonal is normalized to itself, and the lower diagonal is normalized to the maximum of the corresponding ISM reconstruction. The numerical value is the FWHM resolution of the PSF. Each result is shown for increasing saturation factor (ς), from left to right.

We also calculated the so-called fingerprint at increasing STED beam intensities (Fig. 2b). We have defined the fingerprint of an ISM dataset as the sum of all micro-images [34] and it measures the convolution of excitation and detection PSF. Thus, it describes the distribution of photons on the detector array with no influence from the specimen (except for an intensity scale factor). We calculated the fingerprints by integrating along the scanning dimensions (*x_s_, y_s_*) the ISM dataset of the simulated point-source sample. As expected, increasing the STED power reduces the width of fingerprint, reflecting the shrinking of the effective excitation PSF. Eventually, for extreme STED power, the fingerprint identifies with the detection PSF.

To validate our STED-ISM approach, we continued by acquiring superresolved images of various samples. We imaged 20 nm diameter fluorescent beads with increasing STED beam power (Fig. 3a). Consistent with the simulations, for relatively low STED beam power, the STED-ISM image shows better SNR and better resolution, when compared to the conventional STED counterpart (obtained by summing all 25 channels). For high STED beam powers the benefits of STED-ISM become negligible. We confirmed the same results by performing STED-ISM measurements of a sample of fixed Hela cells (Fig. 3b). It is important to note here that the successful STED-ISM reconstruction relies heavily on our APR method, a blind and parameter-free image phase-correlation algorithm [33, 34], able to retrieve the shift-vectors directly from the scanned-images. Although in general this strategy compensates for possible misalignments of the optical system, as discussed above, in the context of STED-ISM it also accommodates implicitly the strong dependency of the shift-vectors on the STED beam powers. In contrast, (i) mechanical ISM implementations should rely on proper modelling or prior calibration of the STED microscope’s effective PSF, as a function of the STED beam intensity and of the general experimental conditions; (ii) optical ISM implementations would require a change of the demagnification factor, which is impractical to achieve.

**Figure 3:**
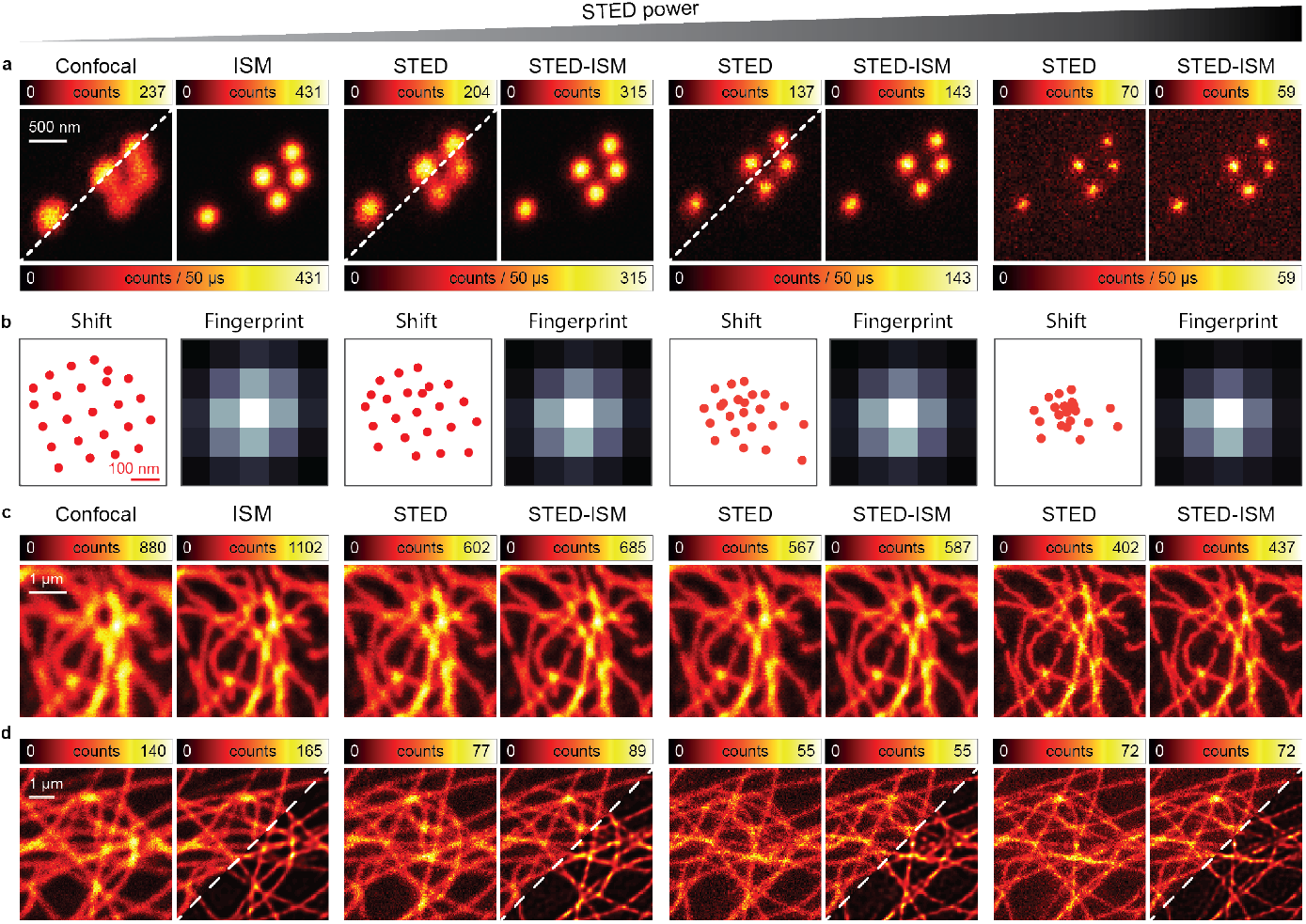
STED-ISM imaging. In **a**, we compare raw images of fluorescent beads with the corresponding ISM reconstructions. In **b**, we show the shift-vectors and fingerprint calculated from an image of the beads. In **c**, we show detailed regions of images of tubulin-labeled fixed cells. In **d**, we show detailed images of living Hela cells with SIR tubulin labeling. More specifically, we compare raw STED (left), STED-ISM (right, upper corner) and the result of multi-image deconvolution STED-ISM^+^ (right, bottom corner). All results are shown with increasing STED power, from left to right. The full images are shown in Suppl. Fig. S3 and S4.

To investigate the concrete advantages of STED-ISM over conventional STED microscopy, we explored the case when one is concerned most about the number of stimulating photons delivered to the sample: live-cell imaging (Fig. 3d, Suppl. Fig. S4). Also in this case, we report an enhancement of SNR and spatial resolution of the resulting STED-ISM images with respect to raw STED counterparts. The results are further improved by applying our multi-image deconvolution algorithm [35, 34], here completely parameter-free thanks to the PSFs estimation *via* Fourier ring correlation (FRC) analysis [36]. Moreover, we were able to perform extended STED-ISM time lapses of live Hela cells without inducing any noticeable photo-bleaching effect, given the reduced STED beam intensity necessary to obtain the target resolution (Suppl. Fig. S5)

### Removing Background with Focus-ISM

In this Section, we regard at the ISM dataset from the perspective of the microimages *i*(***x**_d_*|***x**_s_*) in order to effectively improve the optical-sectioning of STED microscopy by removing the out-of-focus background. In particular, we first introduce our new method, named *focus*-ISM (fISM), in the case of ideal STED, and later we generalise this result to any STED beam intensity. As a matter of fact, our method is well suited also to conventional ISM and any ISM-based technique.

The core idea behind focus-ISM consists in observing the distribution of the light on the detector array to distinguish the axial position of the emitters. We simulate the three-dimensional (3D) scanned PSFs of an ideal STED microscope to demonstrate the working principle (Fig. 4a). The central element of the detector array mostly contains in-focus signal, while the out-of-focus light dominates the outer elements. We can extract the same information by calculating the axial fingerprints (Fig. 4b), namely, by integrating, at different depths *z_s_*, the 3D scanned PSFs over the scanning coordinates (*x_s_,y_s_*). When the emitter is in focus, the central pixels contain most of the signal. The farther the distance from the focal plane, the more the outer pixels of the fingerprint are populated with photons. We can quantify this trend by calculating the ratio of the intensity of the outer pixels to the intensity collected by the central pixel (Suppl. Fig. S6a). As expected, the outer pixels collect more light as depth increases. Interestingly, the micro-images *i*(***x**_d_*|***x**_s_*) of each scan point ***x**_s_* show a similar behaviour (Supplementary Note 2, Suppl. Fig. S7a): in the case of ideal STED, both the micro-images and the fingerprint coincide with the detection PSF, weighted by the sample brightness. Indeed, if the effective excitation spot (the excitation PSF) is small enough to ideally excite only a single point, the corresponding wide-field micro-image is the detection PSF of the microscope, centred at the scan point. Similarly, the fingerprint is the sum over tha scan of all the micro-images, and contains the same information with a higher SNR. For ideal STED microscopy, confinement of the fluorescent region also occurs outside the focal plane (Suppl. Fig. S8). Thus, the out-of-focus micro-images and fingerprints also correspond to the out-of-focus detection PSF. These observations suggest that the information about the axial position is encoded in the lateral distribution of the light at the detector plane, and can be exploited pixel-by-pixel to remove the out-of-focus background. Indeed, each micro-image can be seen as the linear combination of a narrow in-focus component and a broad out-of-focus component.

**Figure 4:**
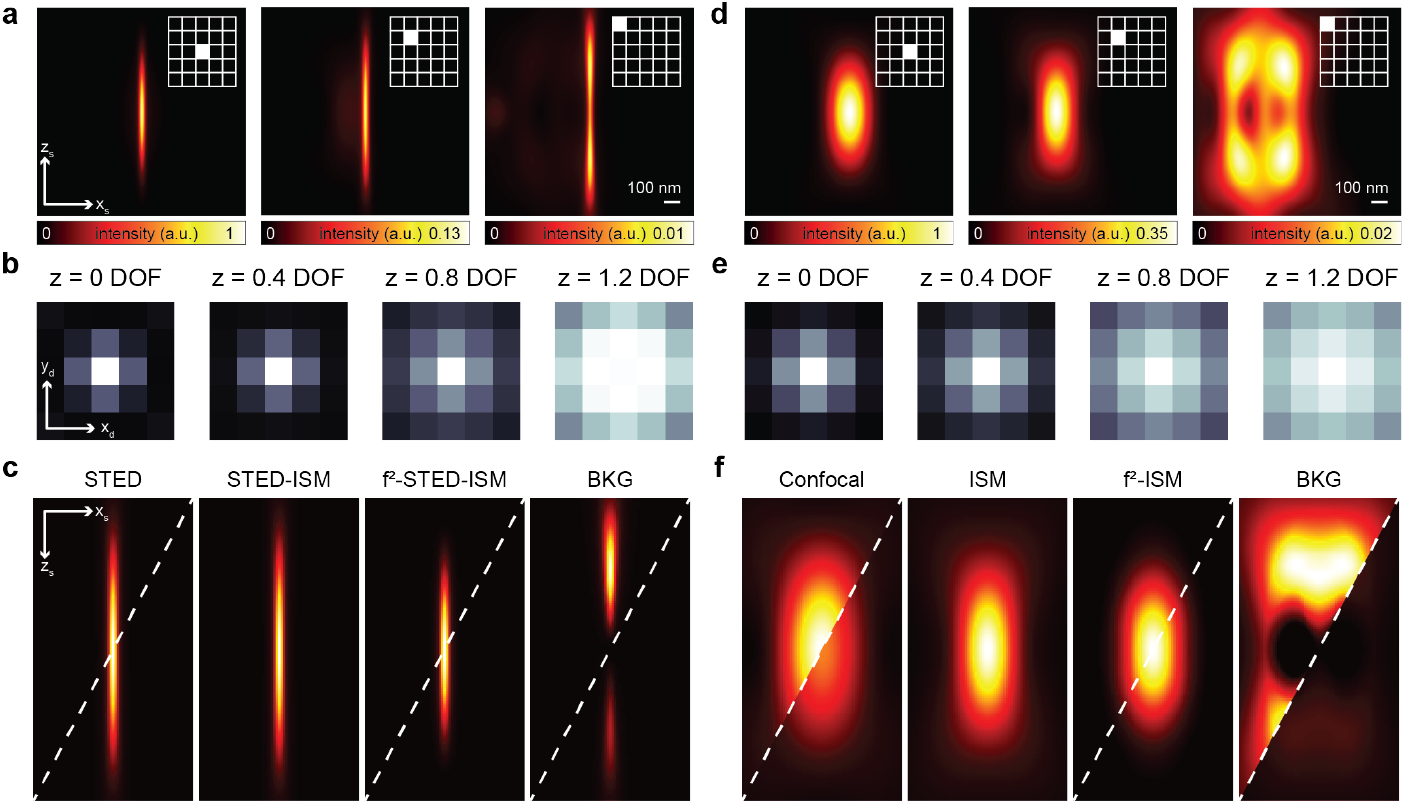
Focus-ISM principle. In **a**, we show STED PSFs (ς = 300) related to the detector element highlighted with the white box. The *z, r* images are obtained by sectioning the 3D PSFs along the main diagonal. In **b**, we show the fingerprint at different axial positions, calculated by summing all the scan points from the corresponding sections of the STED PSF. In **c**, we show the reconstructed STED PSFs. From left to right: raw STED (open pinhole), STED-ISM, focus-ISM, and removed background. In **d**, we show the (normalized) confocal PSFs corresponding to the detector element highlighted with the white box. In **e**, we show the fingerprint at different axial positions, calculated by summing all the scan points from the corresponding sections of the confocal PSF. In **f**, we show the reconstructed confocal PSFs. From left to right: raw confocal (open pinhole), ISM, focus-ISM, and removed background. The top-left corner of each PSF is normalized to itself, while the bottom-right corner is normalized to the ISM reconstruction.

In the following, we present a *naive* approach to exploit the relation between the distribution of the signal on different detector elements and its origin on the optical axis. Later, we discuss a more precise approach to identify and remove the out-of-focus background from each micro-image, and consequently to the reconstructed STED-ISM image.

The out-of-focus light is distributed broadly across the detector elements, and outer elements do not register in-focus light. This observation suggests a simple way to estimate the background – namely using a constant offset. Our first algorithm (*f*^1^MSM) estimates the background by calculating the average of the signal collected by the outer elements of the detector array. The offset is then subtracted from the inner elements. In detail, we use all the outer-ring elements to extract the average background per scan-point. This choice guarantees a conservative estimate of the background. The simplicity behind this strategy, which only requires a few basic operations, enables a fast estimation of the background, paving the way for real-time focus-ISM. However, the approximation behind this approach is crude and might lead to non-physical results, such as pixels with negative photon counts.

Our second approach is more sophisticated and consists in modelling the axial STED fingerprint (i.e., the detection PSF) with a two-dimensional Gaussian distribution, which is known to be a robust approximation [37] in the absence of strong aberrations. We tested the Gaussian model by fitting each simulated axial fingerprint (Suppl. Fig. S6b). As expected, the standard deviation increases for increasing depth. Still, it is approximately constant around the focal plane for roughly one depth-of-field thanks to the 3D structure of the depletion beam (Suppl. Fig. S8). Our second algorithm (*f*^2^-ISM) fits each micro-image to the weighted sum of two Gaussian functions, the first with a broad standard deviation – used to model the out-of-focus component – and the second with a narrow standard deviation – able to model the in-focus light. The standard deviation of the in-focus distribution is kept fixed and precalibrated either theoretically or experimentally (Suppl. Fig. S9). The standard deviation of the out-of-focus distribution is typically a free fitting parameter, but it can also be kept fixed with a user-selectable value (Suppl. Fig. S10). We obtain the final *f*^2^-ISM image by integrating only the portion of the signal fitted to the in-focus term. This second approach is computationally more expensive but enables a more precise classification of the in/out-of-focus components. Additionally, we applied physically meaningful constraints (such as conservation of the photon flux) to guarantee the non-negativity for the pixels of the reconstructed image. Because for ideal STED the shift vectors are null, in this case we did not apply adaptive pixel reassignment before applying focus-ISM.

To demonstrate the capabilities of our algorithm, we applied the *f*^2^-ISM method to the simulated 3D scanned PSF of an ideal STED microscope (Fig. 4c). We recall that in the case of ideal STED the APR method is not necessary. Notably, compared to the STED-ISM PSF, the loss of photons from the focal region is almost negligible, but the out-of-focus light is strongly suppressed, effectively enhancing the optical sectioning of the microscope without reducing the signal. The reliability of our result can also be appreciated by simulating the imaging of a three-dimensional filament network (Suppl. Fig. S11a).

We confirmed the benefit of focus-ISM by performing STED imaging on a sample of tubulin-labeled fixed Hela cells (Fig. 5). This sample is of particular interest because tubulin filaments wrap in three dimensions around the nucleus. In this case, the out-of-focus background significantly degrades the STED-ISM image. The focus-ISM method recovers the contrast significantly by estimating and removing the out-of-focus background. As anticipated, *f*^1^ -STED-ISM may introduce negative intensity values, while this problem vanishes for *f*^2^-ISM. Furthermore, *f*^2^-ISM provides higher signal-to-noise ratio (SNR) than *f*^1^-ISM, as confirmed by the higher peak counts in the image. We further improve image quality with deconvolution: we adapted a multi-image deconvolution algorithm to introduce the background as an *a priori* information (*f*^++^-ISM), which can be estimated with any of the discussed approaches. Notably, using image deconvolution, negative values do not appear, even if the background is estimated using the *f*^1^ -ISM method.

**Figure 5:**
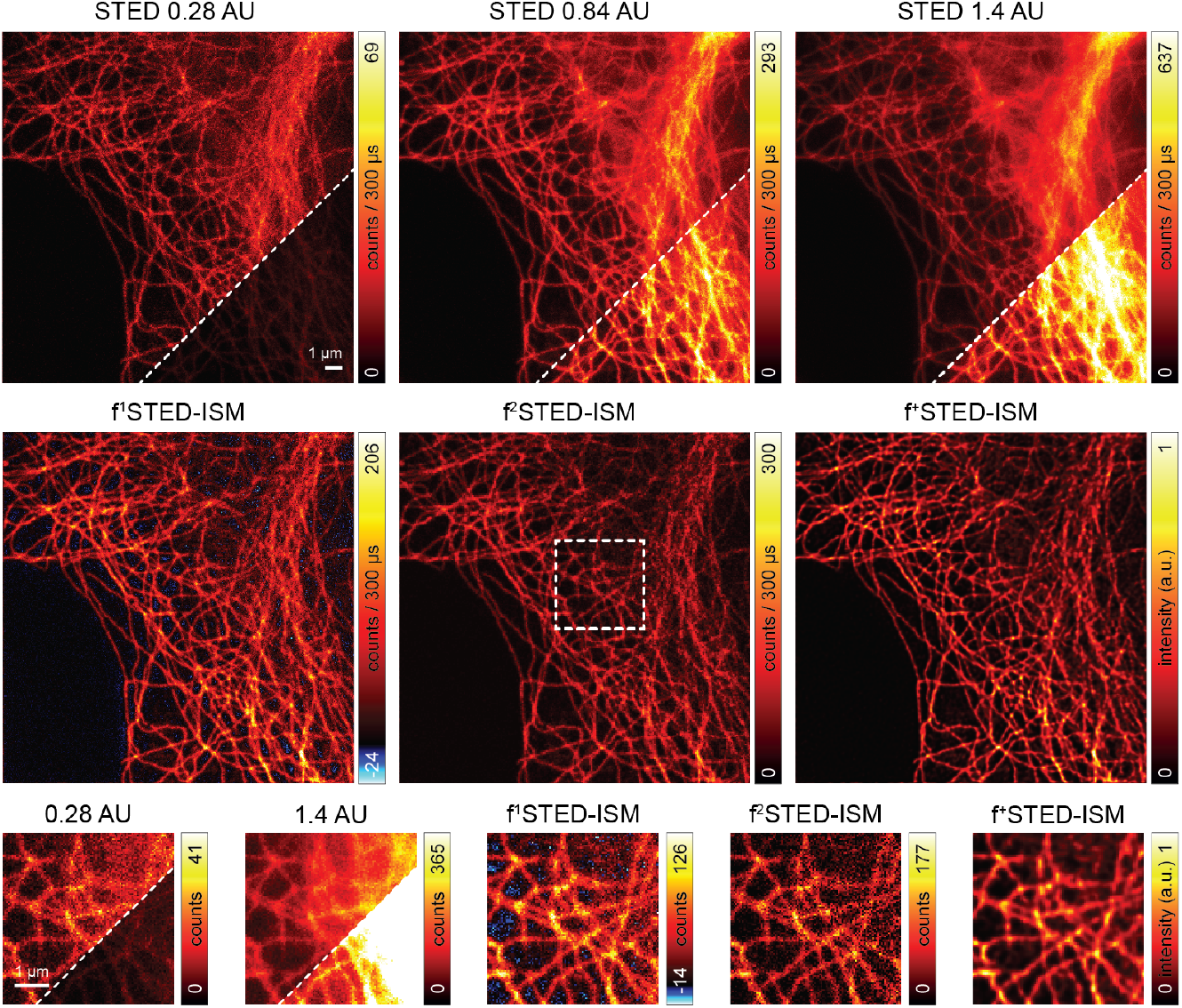
focus-ISM for STED imaging. **Top Row**, STED image of tubulin-labeled fixed Hela cell at different pinhole sizes. The bottom/right corner is normalized to the *f*^2^-STED-ISM image. Even at the smallest pinhole size, the out-of-focus light hides some structure. **Middle Row**, f-ISM reconstruction of the STED image. The *f*^1^ -ISM may introduce some negative values, here colored blue. The *f*^2^-ISM method does not produce negative values and maintains a higher photon count. The *f*^+^-ISM image has been produced with a Richardson-Lucy deconvolution algorithm manually stopped at the 5th step and using the background estimated from the *f*^1^-ISM method. **Bottom Row**, zoomed details of the above images, corresponding to the region identified by the dashed white box.

The advantages of focus-ISM in terms of optical sectioning are even more clear when performing three-dimensional imaging. We show the result of volumetric STED imaging of a tubulin-labeled fixed Hela cell (Fig. 6a, Suppl. Fig. S12). The reconstructed volume, obtained with the *f*^2^-ISM method, presents sharper axial cross-sections and close to no out-of-focus light.

**Figure 6:**
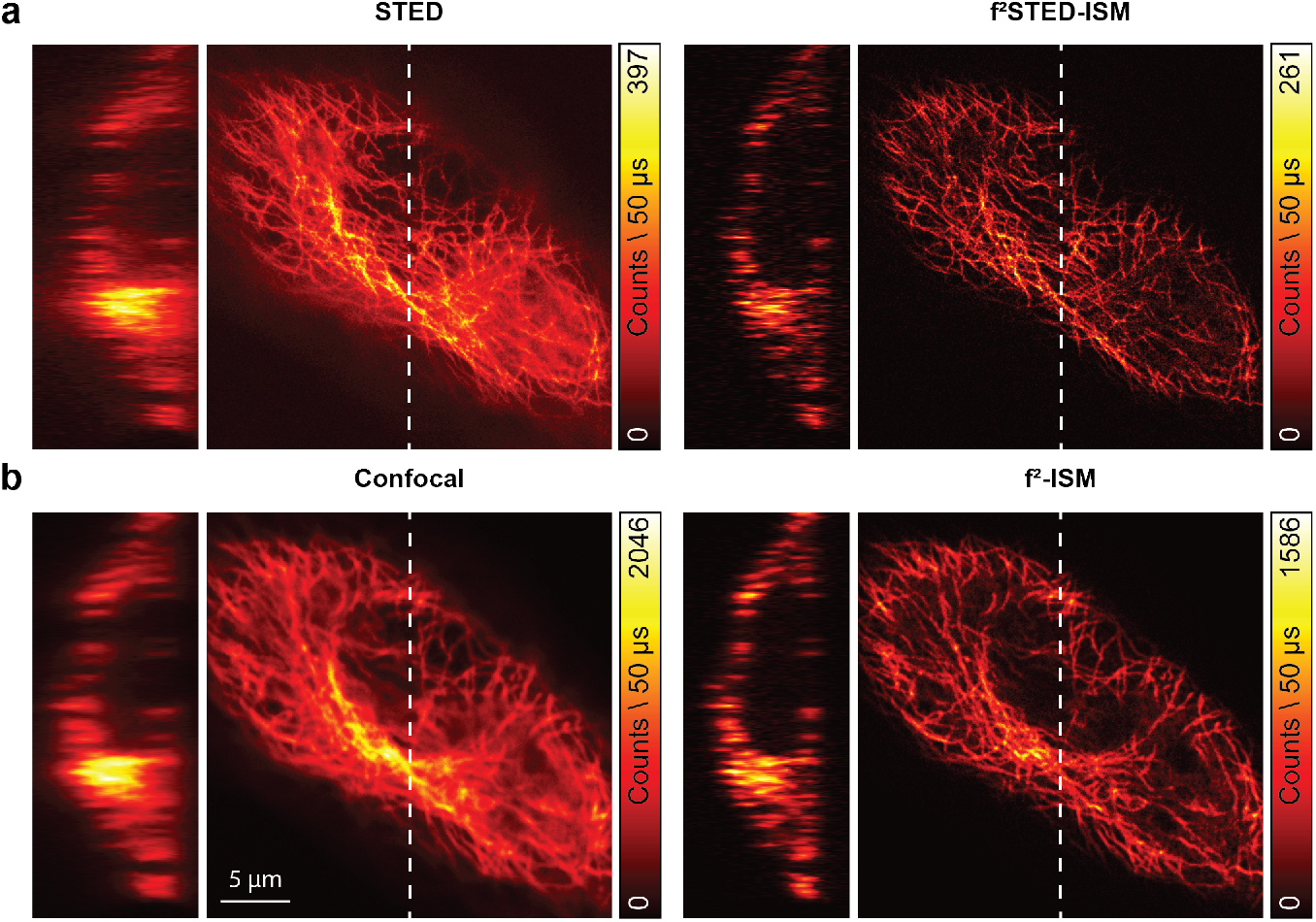
Volumetric focus-ISM. In **a**, we show the STED image of a tubulin-labelled fixed Hela cell (left). The *f*^2^-ISM reconstruction (right) show negligible loss of in-focus photons and greatly enhanced contrast, obtained by removing the out-of-focus content. In **b**, we show the confocal image of the same cell (left). Even in this case, the *f*^2^-ISM reconstruction (right) shows negligible loss of in-focus photons and greatly enhanced contrast. The axial cross-sections are obtained from the x-axis highlighted with a white dashed line.

All focus STED-ISM simulations have assumed only the conventional out-of-focus fluorescence background generated by the excitation beam. However, the anti-Stokes fluorescence background generated by the STED beam is another non-negligible source of background. Because of the annular-shaped distribution of the STED beam intensity, anti-Stokes fluorescence originates mainly from out-of-focus planes or the periphery of the effective excitation region. Thus, the anti-Stokes background localises in the fingerprint and micro-images similarly to the conventional out-of-focus background (Suppl. Fig. S8), and our focus-ISM approach is also able to remove the background from this source.

So far, we have motivated focus-ISM using the hypothesis of high STED beam intensity (i.e., ideal STED microscopy). However, focus-ISM can be applied for any depletion intensity, down to the case of a standard confocal micro-scope. Indeed, we theoretically demonstrate that, as for ideal STED microscopy, the post-APR micro-images are equal to the fingerprint (Supplementary Note 2) for any STED beam intensity. Thus, applying the focus-ISM classification after the APR and then summing the registered scanned images improves the optical-sectioning at any STED beam intensity. An intuitive demonstration of the equivalence between post APR micro-images and fingerprint (Fig. S7c) follows here. Because the micro-images of an ISM dataset are just small wide-field images of the illuminated area (Fig. S7b), their content depends on the structure and the position of the sample. Nevertheless, the result of pixel reassignment is the relocation of each pixel to the position of its emitter on the sample space ***x**_s_*. Consequently, under the hypothesis of perfect reassignment, the pixels of the post APR micro-images carry only the information of the same point ***x**_s_*, as they would under point-like illumination centred at the same position. Thus, the only information content left in the post APR micro-image is the probability of detecting a photon originating from the position ***x**_s_* with the detector element at position ***x**_d_*. This probability distribution is exactly the fingerprint – apart from an intensity scale factor.

Because in the confocal case, the out-of-focus signal is distributed over a broader region than in the ideal STED case (Fig. 4d-e), the detector elements located on the outer ring do not provide a reliable estimate of the background, unless we use a larger detector. Thus, we analysed the confocal images exclusively with the *f*^2^-ISM algorithm. As in the STED case, the confocal PSF reconstructed by focus-ISM contains the same number of photons from the focal region, but the out-of-focus light is almost completely removed (Fig. 4f). We also validated focus-ISM for the confocal case on synthetic images of a threedimensional filament network (Fig. S11b).

We also demonstrated the performance of focus-ISM in the absence of stimulated emission depletion. In detail, we performed confocal imaging of the same tubulin-labeled fixed Hela cell used and we show how focus-ISM removes the background efficiently without sacrificing the SNR (Fig. 6, Suppl. Fig. S13b).

Focus-ISM is drastically different from previous attempts at out-of-focus background reduction. Historically, the first effort to reduce the out-of-focus background consisted in closing a pinhole placed in front of the detector. However, as demonstrated by our results, this approach does not fully solve the problem – some background light can reach the detector even if the pinhole size is small – and compromises the SNR of the resulting image (Suppl. Fig. S14). Three-dimensional deconvolution of a confocal image is a more advanced technique and can, in some cases, be a valuable alternative to focus-ISM. In this approach, by collecting optical sections of a three-dimensional specimen, it is possible to reconstruct the volume by reassigning the photons to their origin. Despite being the most conservative solution – no signal is lost, it is just reassigned – it requires the collection of a whole three-dimensional dataset, forcing much longer acquisition times. In experiments in which it is essential to minimize the light dose (e.g., extreme resolution STED) or when time series of 2D images are the main goal, collecting a full volume image is unfeasible, and so is 3D deconvolution. Thus, a method capable of estimating the out-of-focus background directly from a bi-dimensional image is essential. Other than focus-ISM, some other techniques have been proposed in the past. Subtraction-based methods have been circulating since the early 90s [38] and more recently revamped with the introduction of Airyscan-like detectors [39]. However, sub-traction methods cannot generate spatial frequencies beyond the confocal cut-off frequency, meaning that these methods generate a contrast enhancement rather than an effective optical section. At the same time, the subtraction may introduce negative intensity values, which have no meaning from a physical point of view. The simplest escape route is to assign zero counts to the pixels with negative values, but that would break the linearity property of the image, thus precluding the possibility to associate it with a PSF. To mitigate the introduction of non-physical values, some authors scaled the subtraction term with an empirical factor [40], creating a trade-off between artefact generation and background suppression.

We fully solve these problems with the *f*^2^-ISM implementation of focus-ISM, which can remove the background of a 2D image without sacrificing its SNR and with no risk of generating non-physical results. Indeed, the intensity profiles of the PSFs (Suppl. Fig. S15) confirms that the peak value in focus is negligibly affected. At the same time, the axial extension is significantly smaller. Nevertheless, a critical analysis of focus-ISM is still required. Indeed, the algorithm relies on the assumption that pixel reassignment is exact. However, this condition is verified only if the scanned images are all identical but shifted and rescaled. However, in a real-case scenario, this is only approximately true. The shape of the PSFs associated with each pixel of the detector array is roughly the same for elements inside a region of about one Airy Unit. The PSF of more external elements has a distorted shape, but also carries minimal signal. Thus, the perfect reassignment hypothesis is very robust in the absence of other sources of distortions – such as optical aberrations.

## Discussions and Conclusions

The combination of STED microscopy with a detector array offers impressive advantages in terms of optical sectioning and minimization of the risk of inducing photodamage, key factors for super-resolution imaging of thick and live samples.

As our results demonstrate, the boost in resolution provided by APR and STED synergically combine to enable super-resolution at lower STED beam power. Notably, this benefit is maximal at low saturation factor. Indeed, at higher depletion power the gain in resolution is dominated by the STED effect – ideally unlimited – while the gain in resolution provided by ISM is bounded to a factor of two.

Additionally, the detector array enables the application of focus-ISM, a new algorithm that is capable of discriminating the out-of-focus light from the infocus light. By analysing the post-APR micro-image, we are able to remove the background without introducing artefacts and while preserving the SNR. Thus, the combination of APR with focus-ISM enables high contrast and high-resolution imaging, opening new scenarios for the imaging of thick samples. Remarkably, the proposed method works for any STED beam intensity – including the limiting case of an inactive STED beam – thus offering a new tool also for conventional laser-scanning microscopy equipped with a detector array (e.g. confocal, or two-photon excitation). Importantly, we believe that focus-ISM is just the first step towards a more refined usage of the extra-spatial information provided by the SPAD array. Indeed, in this work we used the axial classification of the light only to remove the out-of-focus light. Nevertheless, we believe that using a more detailed mathematical model it is possible to localize precisely the position of the emitter along the optical axis – similarly to singlemolecule localisation microscopy – paving the way for single-frame multi-plane imaging.

For the sake of completeness, other groups have already used the ISM principle in the context of STED microscopy. In particular, to improve the performance of tomographic STED microscopy [41], and to sustain higher fluorescent photon-flux [42]. However, both these works adopted sub-optimal detector arrays: the former a slow conventional camera and the latter an AiryScan-like detector. While the AiryScan detector – a hexagonally-arranged fibre bundle coupled to a series of photomultipliers (PMT) or single-element SPAD detectors – had the merit of massively spreading the ISM technique in its original computational ISM version, SPAD array detectors have everything in favour to make the definitive transition of laser-scanning microscopy, in general, to detector arrays: SPAD array detectors offer higher elements/pixels scalability, robustness, and higher photon-collection efficiency compared to those of PMTs-based AiryScan. Furthermore, in stark contrast to PMTs, they are single-photon timing detectors.

In this scenario, our STED-ISM implementation based on a SPAD array detector can become the gold standard: the proposed implementation (i) requires only minimal changes in a conventional STED microscopy architecture; (ii) preserves all functions of STED microscopy, such as multi-colour, threedimensional, and fast imaging; (iii) is fully compatible with all current approaches for photo-damage reduction and signal-to-background ratio improvement. In terms of this last point, the single-photon timing nature of the SPAD array detector allows the combination of STED-ISM microscopy and time-resolved STED microscopy [43, 11, 13] to further improve the resolution for a given STED beam intensity. Such a time-resolved STED implementation based on a SPAD array detector will provide benefits not only for imaging but also for fluorescence fluctuation spectroscopy (FFS) [44, 45]: we have recently shown how the SPAD array detector improves the information context of a FFS experiment [46].

In general, this work represents a further fundamental milestone toward the transition from single-element detectors to SPAD array detectors in laserscanning microscopy. The unique ability of a SPAD array to spatially and temporally tag fluorescence photons from the probing volume of a laser-scanning microscope can not only to improve the characteristics of current advanced microscopy techniques, but also open the way to novel techniques.

## Methods

### Custom setup

For this work, we updated the ISM setup described previously [34] with a STED line (Suppl. Fig. S1). Briefly, the excitation beam was provided by a triggerable pulsed (~ 80 ps pulse-width) diode laser (LDH-D-C-640, Picoquant) emitting at 640 nm. The STED beam was provided by a femtosecond mode-locked Ti:Sapphire laser (Chameleon Ultra II, Coherent) running at 775 nm. We coupled the STED laser beam into a 100 m long polarization maintaining fiber (PMF). Before injection into the PMF, the beam passed through two 20 cm long glass rods to temporally stretch the pulse-width to a few picoseconds in order to avoid unwanted nonlinear effects and damage during the fiber coupling. We used a half-wave plate (HWP) to adjust beam polarization parallel to the fast axis of the PMF. The combination of glass rods and PMF stretched the pulses of the STED beam to ≈ 200 ps. We controlled the power of the STED and excitation beams thanks to two acousto-optic modulators (AOM, MCQ80-A1,5-IR and MT80-A1-VIS, respectively, AAopto-electronic). The Ti:Sapphire laser (master) runs at 80 MHz and provides an electronic reference signal which we used to synchronize electronically the excitation laser diode (slave). We used a picosecond electronic delayer (Picosecond Delayer, Micro Photon Devices) to temporally align the excitation pulses with respect to the depletion pulses. The STED beam emerging from the PMF was collimated, filtered in polarization by a rotating GlanThompson polarizing prism and phase-engineered though a polymeric mask imprinting a 0 – 2*π* helical phase-ramp (VPP-1a, RPC Photonics). We rotated a quarter-wave plate and a half-wave-plate to obtain circular polarization of the STED beam at the back-aperture of the objective lens. We co-aligned the excitation and STED beam using two dichroic mirrors (T750SPXRXT and H643LPXR, AHF Analysentechnik). After combination, the excitation and STED beams were deflected by two galvanometric scanning mirrors (6215HM40B, CT Cambridge Technology) and directed toward the objective lens (CFI Plan Apo VC 60 ×, 1.4 NA, Oil, Nikon) by the same set of scan and tube lenses used in a commercial scanning microscope (Confocal C2, Nikon). The fluorescence light was collected by the same objective lens, descanned, and passed through the multi-band dichroic mirror as well as through a fluorescence band pass filter (685/70 nm, AHF Analysentechnik). A 300 mm aspheric lens (Thorlabs) focuses the fluorescence light into the pinhole plane generating a conjugated image plane with a magnification of 300× . For ISM measurements the pinhole is maintained completely open. A telescope system, built using two aspheric lenses of 100 mm and 150 mm focal length (Thorlabs), conjugates the SPAD array with the pinhole and provides an extra magnification factor. The final magnification on the SPAD array plane is 450×, thus the size of the SPAD array projected on the specimen is ~ 1.4 A.U. (at the far-red emission wavelength, i.e. 650 nm). Every photon detected by any of the 25 elements of the SPAD array generates a TTL signal that is delivered through a dedicated channel (one channel for each sensitive element of the detector) to an FPGA-based data-acquisition card (NI USB-7856R from National Instruments), which is controlled by our own data acquisition/visualization/processing software, carma. The software-package carma also controls the entire microscope devices needed during the image acquisition, such as the galvanometric mirrors, the axial piezo stage, and the acousto-optic modulators (AOMs), and visualizes the data. In particular, carma synchronizes the galvanometric mirrors with the photon detection to distribute photons between the different pixels of the images. All power values reported for this setup refer to the power measured before the pair of galvanometric mirrors.

### Sample preparation

We demonstrated the enhancement in spatial resolution obtained with our STED-ISM approach on two-dimensional (2D) imaging of fluorescent beads and tubulin filaments. *Fluorescent beads*. In this study we used a commercial sample of ATTO647N fluorescent beads with a diameter of 23 nm (Gatta-BeadsR, GattaQuant). *Tubulin filaments imaging in fixed cells*. Human HeLa cells were fixed with ice methanol, 20 minutes at −20 °C and then washed three times for 15 minutes in PBS. After 1 hour at room temperature, the cells were treated in a solution of 3% bovine serum albumin (BSA) and 0.1% Triton in PBS (blocking buffer). The cells were then incubated with the monoclonal mouse anti-*α*-tubulin antiserum (Sigma Aldrich) diluted in a blocking buffer (1:800) for 1 hour at room temperature. The *α*-tubulin antibody was revealed using Abberior STAR Red goat anti-mouse (Abberior) for the custom microscope or Alexa 546 goat anti-mouse (Sigma Aldrich) for the Nikon-based microscope. The cells were rinsed three times in PBS for 5 minutes. *SiR-Tubulin in live-cells*. To label tubulin proteins, Human HeLa cell were incubated with SiR-tubululin kit (Spirochrome) diluted in LICS at a concentration of 1 μM for 30 minutes at 37 °C and immediately after imaged in the microscope.

### Numerical simulations

We simulated the point-spread-function of the STED-ISM system using the mathematical model presented in the supplementary information. In more detail, we used a discretized version of the image formation model using the integer indices (*n, m*) ∈ [−2, 2]^2^ to denote the detector element. We applied the adaptive pixel-reassignment method to obtain the PSF of ISM and STED-SIM. Alternatively, we summed all scanned image PSFs to obtain the open pinhole PSF. In this case, the size of the whole array active area represents the size of the pinhole. To calculate the normalised intensity distribution of the excitation PSF, the emission PSF, and the vortex beam we used the Focus Field Calculator Matlab package [47].

Here we list all the parameters chosen for the simulations shown throughout the manuscript. The excitation, emission and depletion wavelengths are, respectively, λ_exc_ = 646 nm, A_det_ = 669 nm and λ_STED_ = 775 nm. All PSFs are calculated in a volume of 1.27 μm × 1.27 μm × 1.27 μm with 127 × 127 × 127 voxels (voxel size = 10nm × 10nm × 10nm). We set the numerical aperture of the oil objective lens to NA = 1.4. We simulated a detector with 5 × 5 sensitive elements, arranged in a squared fashion, with a fill factor of 100% (we neglected any dead area between sensitive elements). We fixed the side length of the detector to 1.4 Airy units (defined by the diameter of the Airy disc, 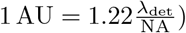). For the STED PSF simulation, we used the following parameters: 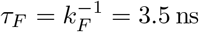, *T* = 1 ns, ς = 0,10, 30, 300, where ς is calculated with respect to the maximum intensity value of the doughnut beam.

### Image reconstruction and analysis

To reconstruct the high-resolution STED-ISM image we used either the simple adaptive pixel-reassignment (APR) method or a multi-image deconvolution algorithm, which are fully described in Castello et al. [34]. Here, we briefly review the two methods.

The adaptive pixel-reassignment (APR) method consists of (i) shifting each scanned-image from detector element (*n, m*) by a shift-vector; (ii) adding up all the shifted images. In this work the shift-vectors are directly estimated from the scanned-images, without the need for any input from the user. In particular, we use a phase correlation approach [33, 48] capable of automatically taking into consideration the geometry of the detector array and the magnification of the microscope system, which can compensate for distortions (misalignments and aberrations) of the system that may arise during imaging. Very importantly in the context of this work, this automatic estimation of the shift-vectors accounts for the saturation level of the STED experiment, i.e. how much the effective fluorescence volume is shrunk, without the need for laborious calibration procedures. Multi-image deconvolution is routinely used when it is necessary to fuse different microscopy images of the same sample, but characterized by different point spread functions [49, 50, 35, 51, 52]. Here, we used the multi-image generalization of the well-known Richardson-Lucy algorithm already introduced by Castello et al. [34]

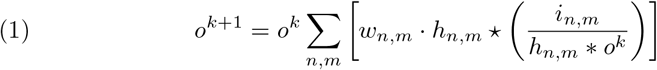

where ⋆ is the cross-correlation operator, * is the convolution operator, *h_n,m_* is the normalized PSF linked to the element (*n, m*) of the SPAD array, *i_n,m_* is the scanned-image generated by the same element, and *o^k^* is the reconstructed image at iteration *k*. The weight factor *w_n,m_*. takes into account the different SNR of the scanned-images and is calculated as the inverse of the fingerprint, i.e. 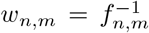. We used a simple Gaussian PSF, shifted by the quantity calculated *via* the phase correlation method. The full width at half maximum (FWHM) of the PSF is fixed by the resolution value calculated by the Fourier ring correlation (FRC) algorithm applied to the STED-ISM image [36]. The same value is used as FWHM for all the PSFs. This protocol results in a sort of blind reconstruction, where no input from the user is required.

In addition, we introduced a small modification to the algorithm to take into account the term *b_n_,_m_*, which is the expected background. In this case, the iterative formula is

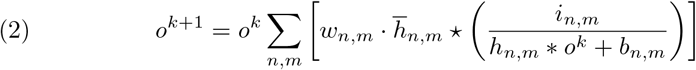

where 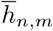 is the (*n, m*) normalized PSF, but not shifted. The PSFs can be measured experimentally and shifted back using the adaptive pixel reassignment method. However, we simplified the algorithm as described previously. Namely, we generated a centred Gaussian PSF with FWHM obtained using the FRC analysis. The background *b_n,m_* can be obtained with any of the focus-ISM implementations described in the following section. Notably, the background term can include also information about the dark-noise. The expected dark noise for each element can be easily measured by registering the signal from the SPAD array detector in the absence of any source of light.

### Focus-STED algorithm

We implemented two versions of our background removal algorithm. The first version, named *f*^1^ -ISM, is the simplest, the fastest, but also the least accurate.

It consists of evaluating pixel by pixel the background *β* as the average intensity value of the external frame of the 5 × 5 micro-image

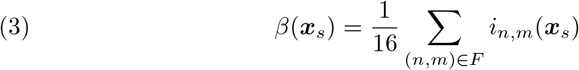

where *F* = {(*n, m*) : |*n*| = 2 ∨ |*m*| = 2}. Then, the in-focus signal is estimated as

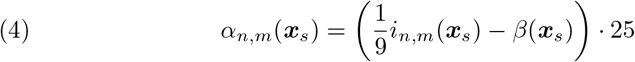

If, as a result, some pixels have negative values, they are trimmed to zero. The second version, named *f*^2^-ISM, is more computationally demanding but is more accurate and cannot generate non-physical results. First, a region of the image containing only in-focus emitters is manually selected to calculate the in-focus fingerprint. The latter is fitted to a single Gaussian function and its standard deviation is recorded as *σ*_sig_. If it is not possible to identify a region that contains only in-focus emitters, then an additional calibration measurement is needed. Alternatively, it is possible to estimate *σ*_det_ theoretically. Then the adaptive pixel reassignment method is applied to the full dataset. Subsequently, each reassigned micro-image of each pixel is normalized and fitted to the following model

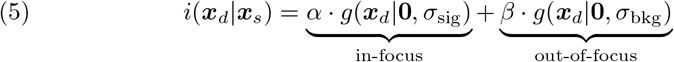

where *g*(***x***|***μ***, *σ*) is a normalized Gaussian function of average ***μ*** and standard deviation *σ*. The weights *α* and *β* follow the conservation of photon flux constraint, namely *α* + *β* =1. The standard deviation of the background micro-image *σ*_bkg_ can be either selected manually or left as a free fitting parameter (see Supp. Fig. S10). In the latter case, only two parameters are free. However, individual micro-images typically have a very poor SNR. Therefore, it is good practice to restrict the minimum value of σ_bkg_ to avoid overfitting. The parameters found with the fitting algorithm are used to build two different micro-images for each pixel: the two Gaussian functions generate, respectively, an in-focus and an out-of-focus micro-image. Eventually, the pixels of each classified micro-image are summed to generate the in-focus and out-of-focus image.

## Acknowledgements

This research was supported by Fondazione San Paolo Observation of bio-molecular processes in live-cell with nanocamera no. EPFD0098 (to G.V.), and by the European Research Council, BrightEyes no. 818699 (to G.T., and G.V.). We thank Prof. Alberto Tosi, Dr. Mauro Buttafava, and Federica Villa – from Politecnico di Milano – for the joint development of the SPAD array detector and for the continuous support in its optimisation; Sabrina Zappone, Andrea Bucci, Marco Scotto, Dr. Mattia Donato, Dr. Eli Slenders and Dr. Eleonora Perego – from Istituto Italiano di Tecnologia – for useful discussions; Elena Tcarenkova – from University of Turku – for helpful discussions in the early stages of this project. Dr. Luca Lanzano, Dr. Michele Oneto, and Simone Pelicci – from Istituto Italiano di Tecnologia – for support in sample preparation.

## Author contributions

G.T., A.Z., and G.V. designed the study. G.V. supervised the project with support from C.J.R.S, P.B., and A.D. G.T., F.F., S.K., A.Z., and G.V. designed and implemented the custom STED system with the SPAD array detector. G.T., and M.C. developed the control software. S.P., and M.C. integrated the SPAD array detector in the control and data-acquisition system. G.T., A.Z., M.C., S.K. and G.V., developed the analysis software. G.T., F.F., A.Z., and G.V., performed the experiments. G.T., A.Z., and G.V. analyzed the data with the support of all other authors. G.T., A.Z., and G.V. wrote the manuscript. All authors discussed the results and commented on the manuscript.

## Supplementary Material

### 1 Adaptive Pixel Reassignment

From a statistical point of view, the pixel-reassignment (PR) concept reassigns the micro-image’s photons *i*(***x**_d_*|***x**_s_*) to the most likely position of origin in the sample space ***x*** (Fig. 1b). For a given scanning position ***x**_s_*, the most likely point from which the photons detected at the position ***x**_d_* originate is the maximum point (mode) of the joint probability, or effective point-spread function, *h* = *h*_exc_ × *h*_det_. Here *h*_exc_ is the excitation PSF, i.e., the probability to excite a fluorophore as a function of its position, and *h*_det_ is the detection PSF, i.e., the probability of detecting a photon – at ***x**_d_* – as a function of the fluorophore position. If we denote with ***μ***′(***x**_d_*) the displacement between the detection position ***x**_d_* and the effective PSF maximum – known as the micro-image shift-vector, PR consists in repositioning the photon location from ***x**_d_* to ***x*** = ***x**_s_* – ***x**_d_*+***μ***′(***x**_d_*). Notably, if ***μ***′(***x**_d_*) is null, the resulting PR image is the conventional wide-field image. Instead, if ***μ***′(***x**_d_*) = ***x**_d_* then all photons are assigned to the scanning position ***x**_s_* and the PR image is the conventional LSM image (with an open pinhole) [1].Given the micro-images and the relative shift-vectors, the PR operation can be easily implemented computationally.

To avoid post-processing, different opto-mechanical PR implementations have been proposed, assuming no Stokes-shift, and Gaussian shaped excitation and detection PSFs. Under these assumptions, the maximum of the joint probability is located at the middle of the detection and excitation PSFs’ positions, namely ***μ***′(***x**_d_*) = ***x**_d_*/2. Mechanical ISM [2, 3] implements PR by re-scanning the fluorescent signal with twice larger scan amplitude – than used by scanning the excitation beam. Optical ISM [4, 5] adopts a multi-spot excitation scheme and implements the PR by demagnifying twice the fluorescent spots. Interestingly, some computational ISM implementations [6, 7] also use the multi-spots scheme to speed-up imaging rate.

Practically, however, the Stokes-shift is not negligible, and the excitation and detection PSFs may deviate from ideal Gaussian shapes. These aspects reduce the versatility and robustness of the opto-mechanical ISM implementations. Computational ISM implementations require strategies to effectively determine the micro-image shift-vector ***μ***′(***x**_d_*). In this context, we have introduced the adaptive pixel-reassignment (APR) method, which automatically adapts to the Stokes-shift and to realistic PSF shapes. To explain the APR method, it is necessary to regard the raw ISM dataset from a different point of view: each element of the detector array can be considered as an independent detector generating a confocal image with an offset pinhole, the so-called scanned-image. In other words, the raw ISM dataset can be regarded both as a collection of micro-images *i*(***x**_d_*|***x**_s_*) – one for each sample position ***x**_s_*, or a collection of scanned-images *i*(***x**_s_*|***x**_d_*) – one for each detector position ***x**_d_*. From the latter point of view, the PR method reassigns the scanned-image’s photons *i*(***x**_s_*|***x**_d_*) to the most likely position of origin in the sample space ***x***. Because this position is encoded in the scanned-image, we obtain the scanned-image shift-vector ***μ***(***x**_d_*) by extracting the relative displacements between the scanned-images. Thus, the new position of the photons reads ***x*** = ***x**_s_* – ***μ***(*x_d_*).

Notably, given the scanned-image shift-vector one can derive the micro-image shift-vector, i.e., ***μ***′(***x**_d_*) = ***x**_d_* – ***μ***(***x**_d_*). The APR is effectively implemented by shifting the scanned images by ***μ***(***x**_d_*) and summing them up.

### 2 Working principle of Focus-ISM

The point-spread function (PSF) of the scanned-image generated by the element at position ***x**_d_* of a detector array is given by the PSF of a laser scanning microscope, just with a shifted pinhole aperture [8, 9]. The equation is

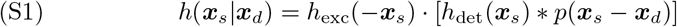

where ***x**_s_* = (*x_s_, y_s_*) are the scan coordinates on the sample plane in focus, *p* is the pinhole aperture that describes the sensitive area of the element of the detector array located at ***x**_d_* = (*x_d_, y_d_*), and *h*_exc_ and *h*_det_ are, respectively, the excitation and detection normalized PSFs 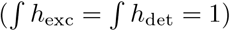.

For STED-ISM, we also have to consider the reduction of the effective fluorescence region, which is equivalent to a reduction of the effective excitation PSF

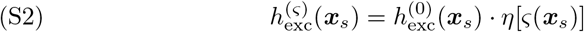

where 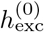 is the confocal excitation PSF, and *η* is the depletion function, which describes the reduction of fluorescence emission under exposure of the depletion beam. ς is the saturation factor, defined as

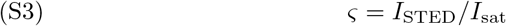

where *I*_STED_(***x**_s_*) is the doughnut-shaped intensity distribution of the STED beam and *I*_sat_ is the saturation intensity, namely the intensity of stimulated emission needed to have spontaneous emission as likely as stimulated emission. In other words, the spontaneous emission rate *k_F_* equals the stimulated emission rate *k_SE_* when *I*_STED_ = *I*_sat_. In order to calculate the saturation factor ς using a pulsed laser, we need to take into account the dynamics of the fluorescence intensity *F*(*t*) under stimulated emission [8]. The fluorescence decay is

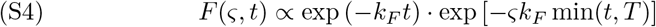

where *T* is the pulse duration of the STED beam. Assuming a time-gated detection in the microscope (namely, the light is collected only after the STED beam pulse is over), the depletion function for the gated pulsed STED implementation reads

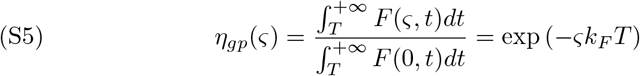

Finally, the equation for the depleted excitation PSF is

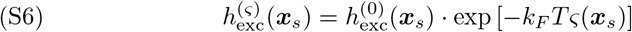

Since ς(***x**_s_*) is doughnut-shaped, the excitation PSF is more depleted at its borders and its effective width is smaller. Notably, the STED excitation PSF is no longer normalized.

For the rest of the manuscript, we will indicate with *h*_exc_ a generic excitation PSF with effective width depending on the saturation factor. Thus, the scanned-image generated by the detector element at position ***x**_d_* can be written as

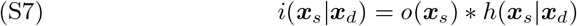

where *o*(***x**_s_*) is the specimen, here considered as a distribution of light emitters. In this work, we consider the limit case where each element of the SPAD array can be described as a shifted Dirac delta function. Substituting *p*(***x**_s_*) with *δ*(***x**_s_* – ***x**_d_*) in equation S1 we obtain

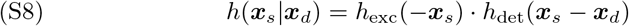

In order to grasp the core idea behind image scanning microscopy (ISM), we first introduce its concept using a Gaussian approximation and later proceed with the general case.

#### 2.1 Gaussian model

In this subsection, we simplify the model approximating the emission and excitation PSF with a Gaussian distribution with circular symmetry,

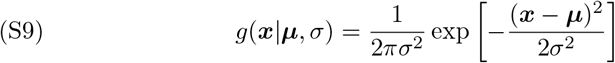

where ***μ*** is the mean and *σ* is the standard deviation. Thus, the resulting PSF is

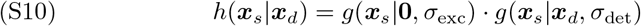

Notably, the product of two Gaussian function is another Gaussian function with the following properties:

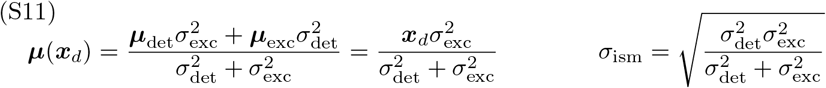

where ***μ***(***x**_d_*) is known as *shift-vector*, and σ_ism_ is independent of ***x**_d_*. The intensity of the resulting Gaussian function is also rescaled by the following rescaling factor

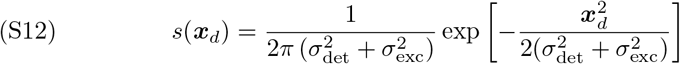

Thus, the resulting PSF can be written as

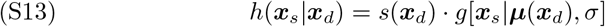

Neglecting the Stokes-shift, we can assume *σ*_exc_ ~ *σ*(_det_ to obtain the well-known 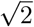 gain in resolution. However, the images generated by each detector element are shifted with respect to the central element. Indeed, the scanned-image is

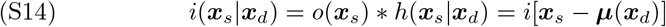

If all the images are simply summed together, the resolution gain is lost, and the resulting image is equivalent to that generated by a traditional confocal microscope with an open pinhole (as large as the detector size). The pixel reassignment (PR) algorithm shifts back each image

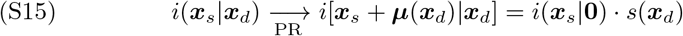

and constructs a new image summing the reassigned images

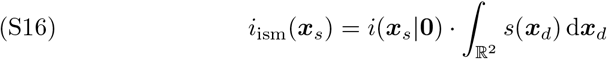

thus conserving the gain in resolution and enhancing the signal-to-noise ratio (SNR).

#### 2.2 General model

The typical assumption behind ISM is that the complete PSF can still be written as a new shifted function

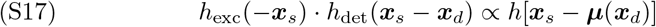

Thus, the scanned-images generated by each detector are all of similar shape, but shifted,

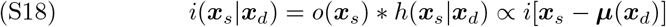

Instead of using the theoretical values found using the Gaussian approximation, the shift vectors are found by the adaptive pixel reassignment (APR) algorithm. This latter first calculates the correlogram

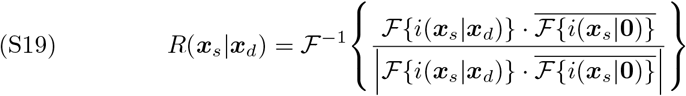

and later finds the shift vectors as the position of maximum correlation

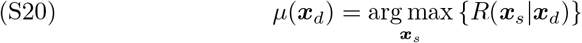

and shifts each scanned image of the corresponding shift vector

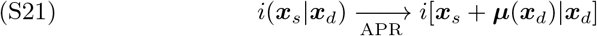

with the purpose of making all the scanned-images identical, except for an intensity scale factor

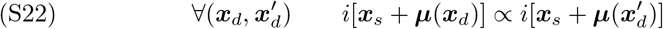

Rather than considering the scanned-image, the ISM image formation process can be seen equivalently from the perspective of the detector. In fact, the detector array can be considered as a small camera, capable of acquiring wide-field images that we call *micro-images*. The equation of these latter can be found as follows:

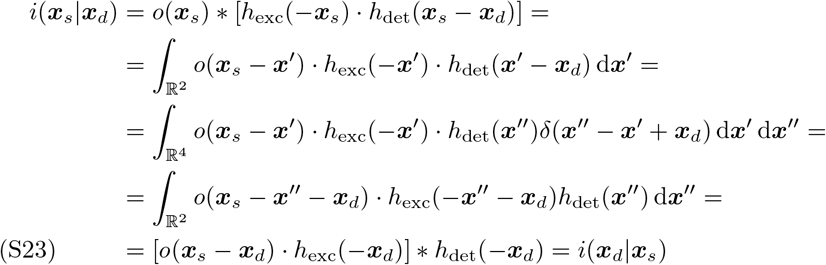

As expected, the micro-image is simply given by the wide-field image formation law, but applied to the illuminated object, i.e. the object multiplied by the excitation PSF. After APR, the reassigned micro-image is

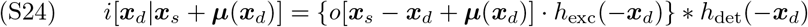

For clarity, we now rewrite the new coordinates of the micro-images as

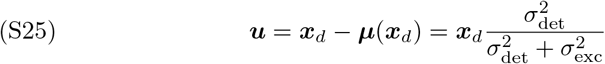

where we have used the theoretical value of the shift-vectors found using the Gaussian approximation for simplicity. In this new coordinate system, the post-APR micro-image is

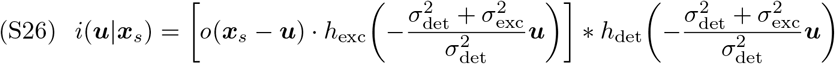

In other words, the effect of APR is a digital shrinking of the excitation and detection PSFs with respect to the object. In detail, the standard deviations of the PSFs become

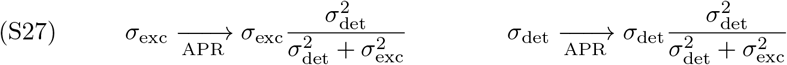

which are always smaller than the original standard deviation.

The relation S22 applies also to the micro-images

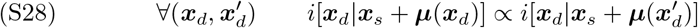

Notably, the proportionality factor in the above relation is independent of ***x**_s_*. Thus, the above condition is satisfied only if equation S24 can be factored into a term depending only on ***x**_s_* and another term depending only on ***x**_d_*. This can occur only in the following cases:

1. If *o* is constant over the whole space, namely it is a flat object. This case is trivial and we will not consider it.
2. If ***μ***(***x**_d_*) = ***x**_d_*, then *o* depends only on ***x**_s_* and is constant with respect to ***x**_d_*. This case is satisfied only if *h*_exc_ = const, namely using a wide-field illumination. This would come at the cost of sacrificing resolution and it is not of our interest.
3. If it is possible to achieve a perfect pixel reassignment. This case is possible only if we can rewrite the product *h*_exc_(–***x**_s_*) · *h_det_*(***x**_s_* – ***x**_d_*) exactly as a new shifted function *h*(***x**_s_* – ***μ***). Namely, the shape of the complete PSF h has to be independent of ***x**_d_*. This condition can be satisfied in ideal confocal microscopy.
4. If the illumination spot is so small that only the position ***x**_s_* of the object o contributes to image formation. This case is satisfied only using a pointlike excitation *h*_exc_(***x**_d_*) = *δ*(***x**_d_*), as in ideal STED microscopy.

We are interested only in the third and fourth case, analyzed in detail in the following sections.

#### 2.3 The confocal case

In this section, we assume that the following relation is exact

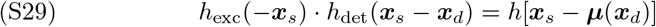

where the complete PSF *h*[***x**_s_* – ***μ***(***x**_d_*)] in general is not normalized. The normalization factor is calculated as

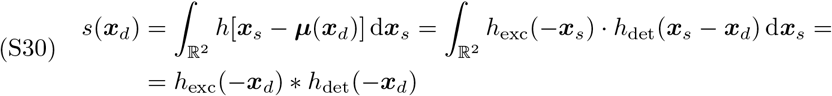

Thus, we can rewrite the PSF as

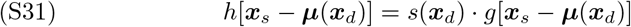

where *g*(***x**_s_*) is a normalized function. Using assumption S29, we replicate the calculations that led to equation S23 after performing APR

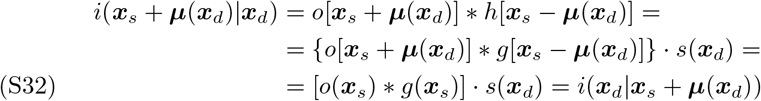

Notably, the scaling factor *s*(***x**_d_*) is exactly the convolution of the excitation PSF with the detection PSF. Thus, we can write the equation of the micro-image after APR as

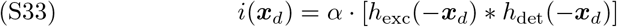

where *α* = *o*(***x**_s_*) * *g*(***x**_s_*) is a proportionality factor with the units of a photon flux. Thus, the reassigned micro-image equals the *fingerprint* of the image, except for an intensity scale factor. The *fingerprint f*(***x**_d_*) of the system is defined as the integral of the image *i*(***x**_s_*|***x**_d_*) over all the scan points ***x**_s_*

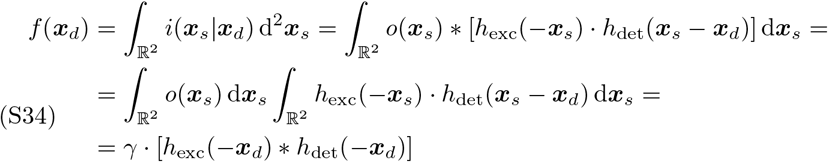

where 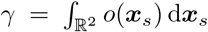 is a scale factor that depends only on the sample distribution.

Therefore, we have found that under the assumption S29, the reassigned micro-image is proportional to the fingerprint. Importantly, this mathematical relation is satisfied by a Gaussian PSF, which is well known to be a good approximation for both detection and excitation PSFs if ||***x**_d_*|| < 1 Airy Unit. However, our result is more general, being valid for each function that satisfies equation S29. Moreover, APR is an adaptive algorithm which can compensate for mild deviations from ideality, since the shift-vectors are calculated as the translations that maximize the similarity between the scanned images.

#### 2.4 The STED case

Assuming an ideal STED system, we can impose a point-like excitation

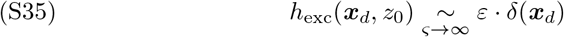

where *ε* is a (theoretically vanishing) scale factor. In this case, the shift vectors are null

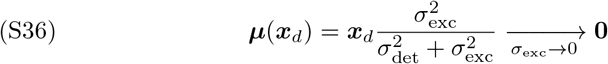

Thus, APR does not modify the micro-images. Using assumption S35, equation S23 becomes

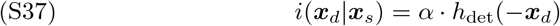

where *α* = *ε* · *o*(***x**_s_*) is a proportionality factor with the units of a photon flux. We found the interesting result that the micro-image obtained while performing STED microscopy is the detection PSF.

Applying condition S35 to equation S34, we obtain the fingerprint of a STED image

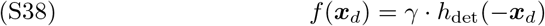

which corresponds to the detection PSF, except for a scale factor.

#### 2.5 Focus-ISM

We have found that, after APR and under reasonable assumptions, the micro-images of an ISM dataset are proportional to the fingerprint of the image. In this section, we generalize the equations for the micro-images and for the fingerprint to the case of a three-dimensional object *o*(***x**_s_, z*). The micro-image of a 3D object is

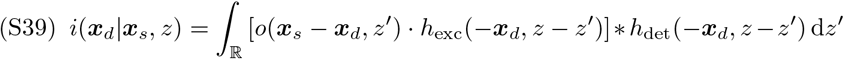

where the convolution is calculated along the dimensions (*x_d_, y_d_*). Assuming that the sample is localized on a single plane, we have *o*(***x**_s_* – ***x**_d_, z*) = *o*(***x**_s_* – ***x**_d_*)*δ*(*z* – *z*_0_). Using this assumption, we obtain the micro-image of an object o placed at distance *z*_0_ from the image plane *z*

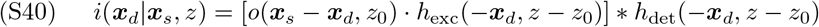

After pixel reassignment, the micro-image becomes

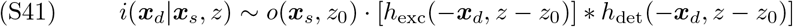

The *fingerprint* of an image generated by a 3D distribution of emitters *o*(***x**_s_, z*) is

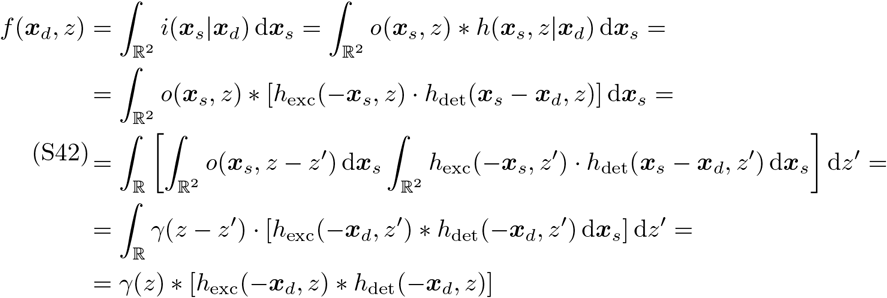

where 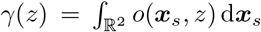 is a weight function that depends only on the sample distribution, and the first convolution is calculated over the *z*-axis, while the second convolution is calculated over the ***x**_s_* axes. Using the assumption that the sample is localised on a single plane, we have *γ*(*z*) = *γ*_0_*δ*(*z* – *z*_0_) and we obtain

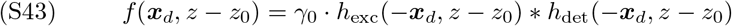

which is proportional to equation S41. We can use this result to write the equation of a micro-image generated by multiple emitters at different axial planes

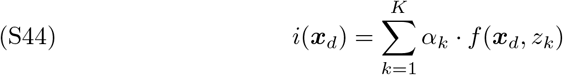

Notably the fingerprint is always centered with respect to the detector array, but its lateral width depends on the axial position of the emitter. Approximating again each PSF with a Gaussian function, the fingerprint is a Gaussian function as well. Considering only two contributes (in-focus and out-of-focus), we finally obtain

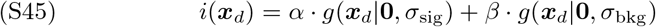

where the weights of the Gaussian functions respect the conservation of the energy constraint

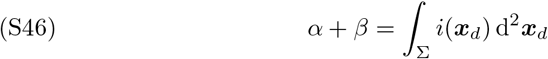

Notably, the standard deviation of the in-focus component can easily be measured experimentally. The fingerprint is effectively a micro-image averaged over many scan points. Thus, the SNR of the fingerprint is much higher than that of a micro-image. We can exploit this information to calibrate *σ*_sig_. This process can be performed with a separate experiment, or alternatively, it is possible to calculate the *fingerprint* on a region of *i*(***x**_s_*|***x**_d_*) containing only in-focus emitters. If fitted to a Gaussian function, the fingerprint of the sub-image provides an experimental measure of *σ*_sig_.

Once each micro-image has been separated in an in-focus and out-of-focus component, the corresponding images are constructed as follows

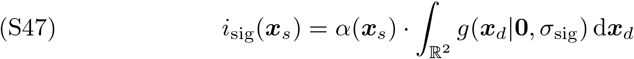

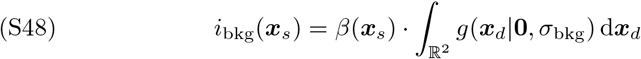

---

**Figure S1:**
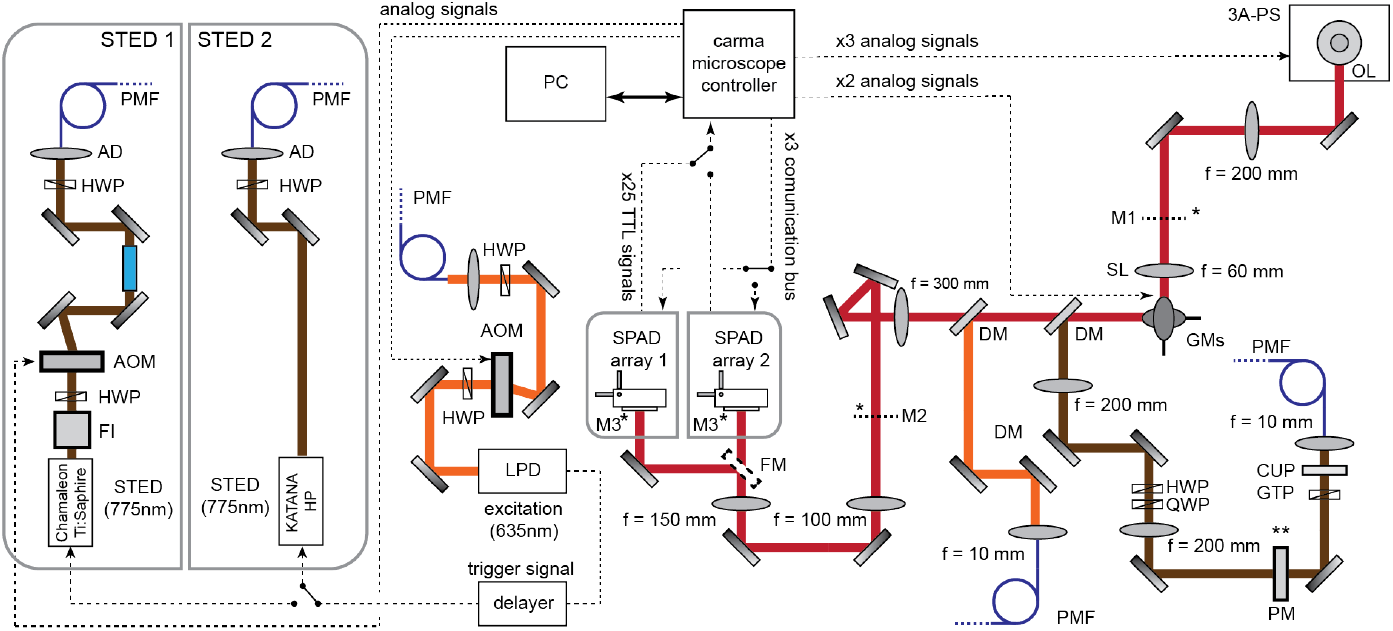
The STED-ISM setup. HWP:half-waveplate;AOM:acousto-optic modulator; AD:achromatic doublet; PMF: polarized-maintaining fiber; FI: Faraday isolator; GR: glass-road; 3A-PS: three-axis piezo stage; SL: scanning lens; GNs: galvanometric mirrors; DM: dichroic mirror; FM: flip mirror; QWP: quarter-wave plate; BPF: band pass filter; MMF: multi-modes fiber; APD: avalanche photo-diode; PM: phase-mask; GTP: Glan-Thompson polarizer; CUP: clean-up filter. Magnifications: M1 = 60×, M2 = 300×, M3 = 450×. The asterisk denotes the plane conjugate to the image plane. The double asterisks denote the plane conjugate to the objective back-aperture.

**Figure S2:**
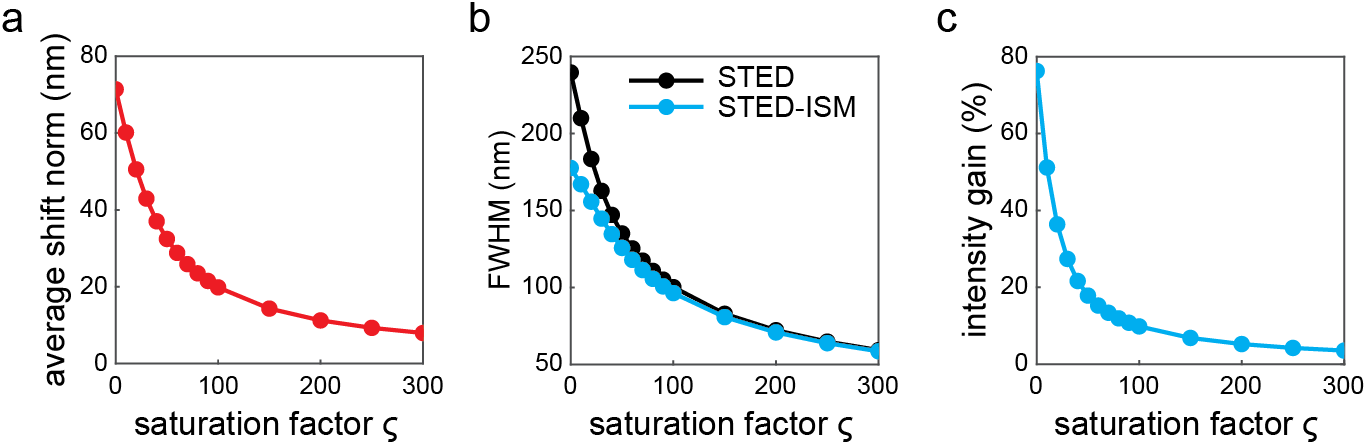
STED-ISM benefits. In **a**, we calculate the average norm of the shift vectors as a function of the saturation factor (ς). For increasing STED power, the images are less shifted. In **b**, we compare the resolution (calculated as the FWHM of the PSF) of raw STED versus STED-ISM. In **c**, we present the reduction of beam power that we can apply to maintain the same lateral resolution of raw STED. The benefit of STED-ISM is maximal at low saturation factors.

**Figure S3:**
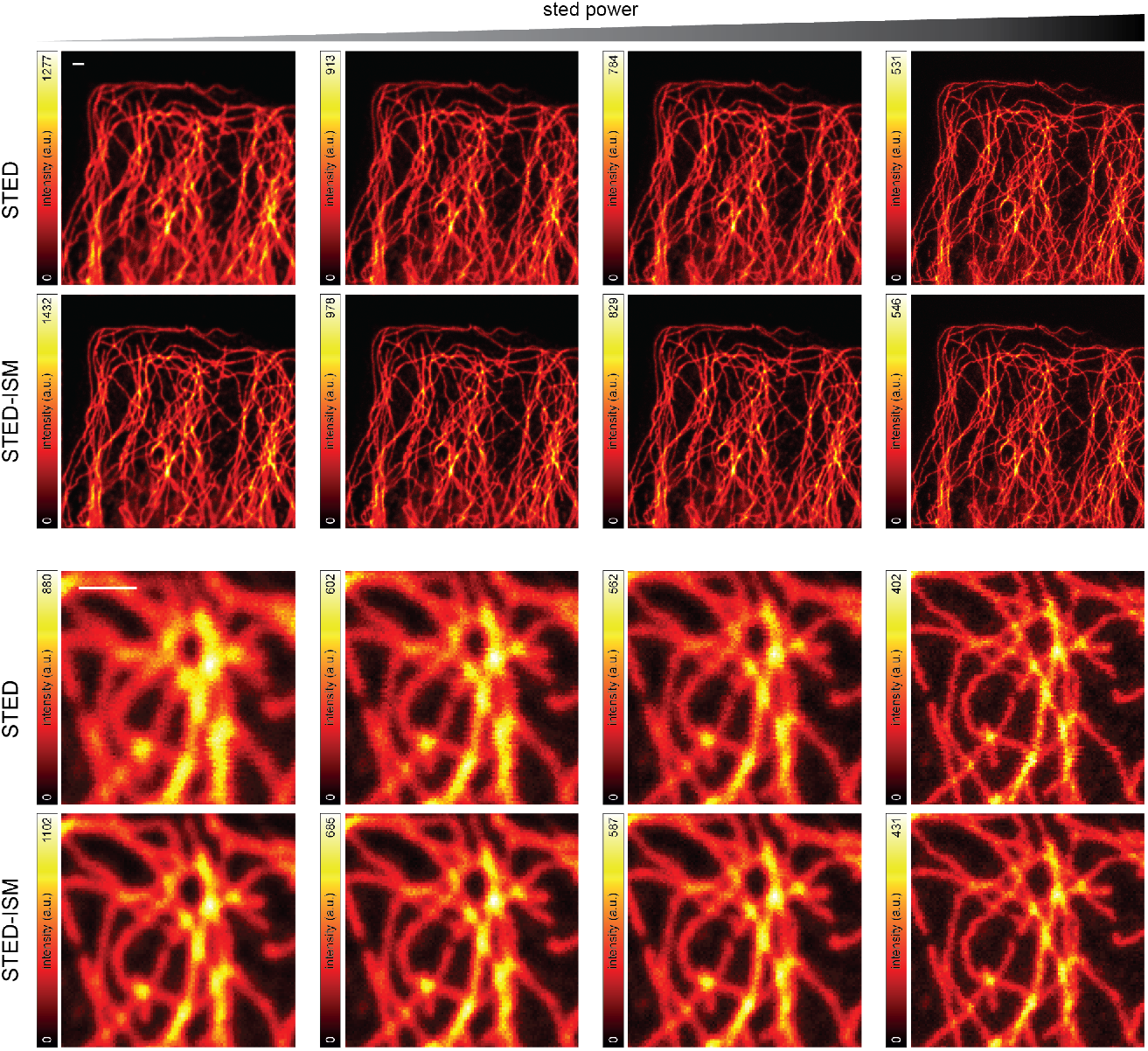
STED-ISM measurements of fixed cells with increasing STED power. The resolution and the SNR of are enhanced by STED-ISM when imaging tubulin-labelled fixed Hela cells. Retrieved FRC resolution values: 278 and 189 nm, 202 and 180 nm, 192 and 177 nm, 136 and 136 nm, for raw STED and STED-ISM, respectively. Dwell time: 100 μs; pixel size: 40 nm; STED powers: 0, 15, 23, 137 mW; format: 500 × 500 pixels; details format: 150 × 150 pixels; all scale-bars: 1 μm.

**Figure S4:**
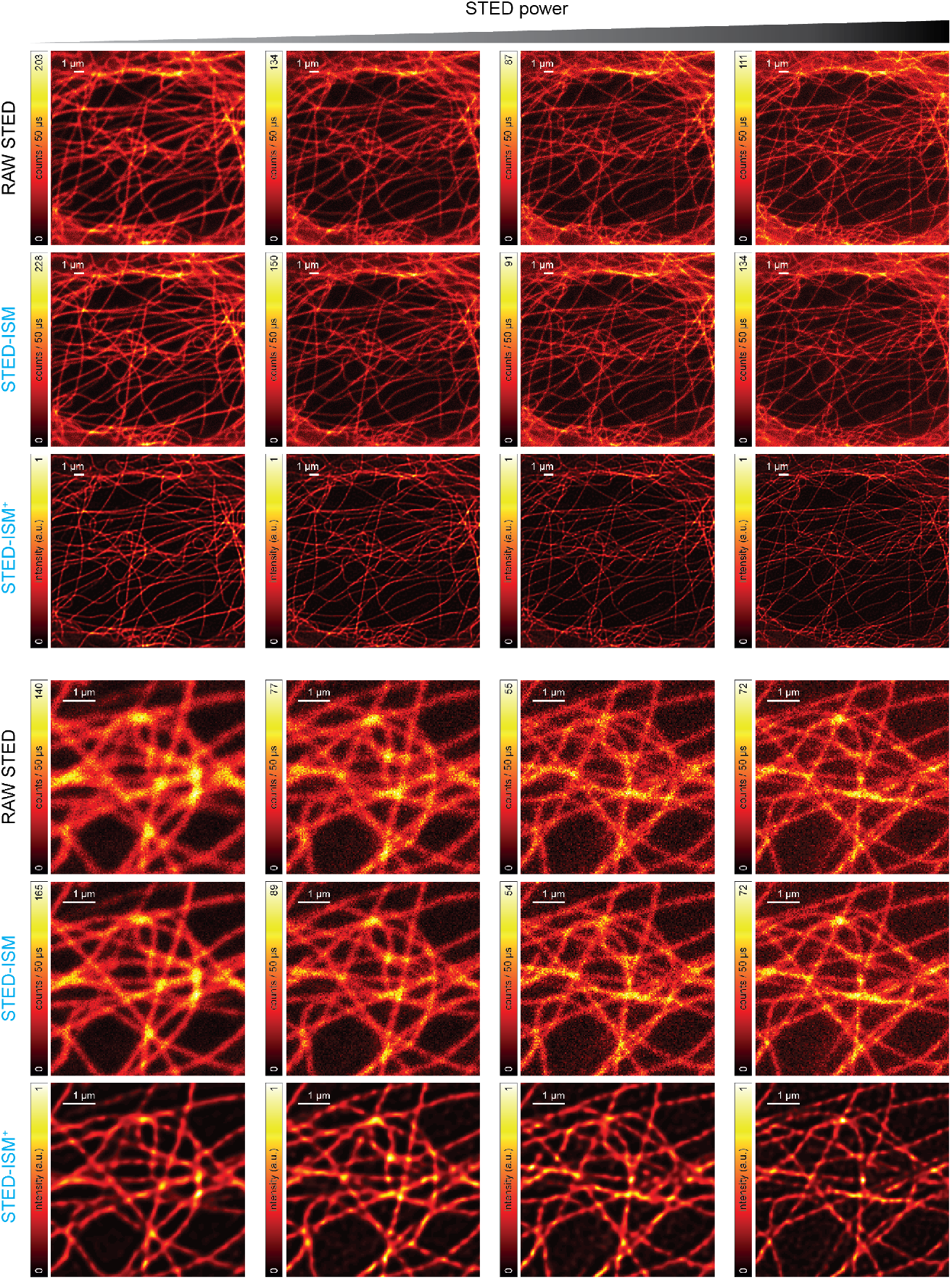
Live cell STED-ISM imaging with increasing STED power. Retrieved FRC resolution values: 290 and 213 nm, 233 and 202 nm, 202 and 202 nm, 168 and 165 nm, for raw STED and STED-ISM, respectively. Pixel dwell time: 50 μs. Pixel size: 40 nm. STED powers: 0, 10, 20, 45 mW. Format of the full images: 500 × 500 pixels. Format of the details: 150 × 150 pixels. All scale-bars: 1 μm

**Figure S5:**
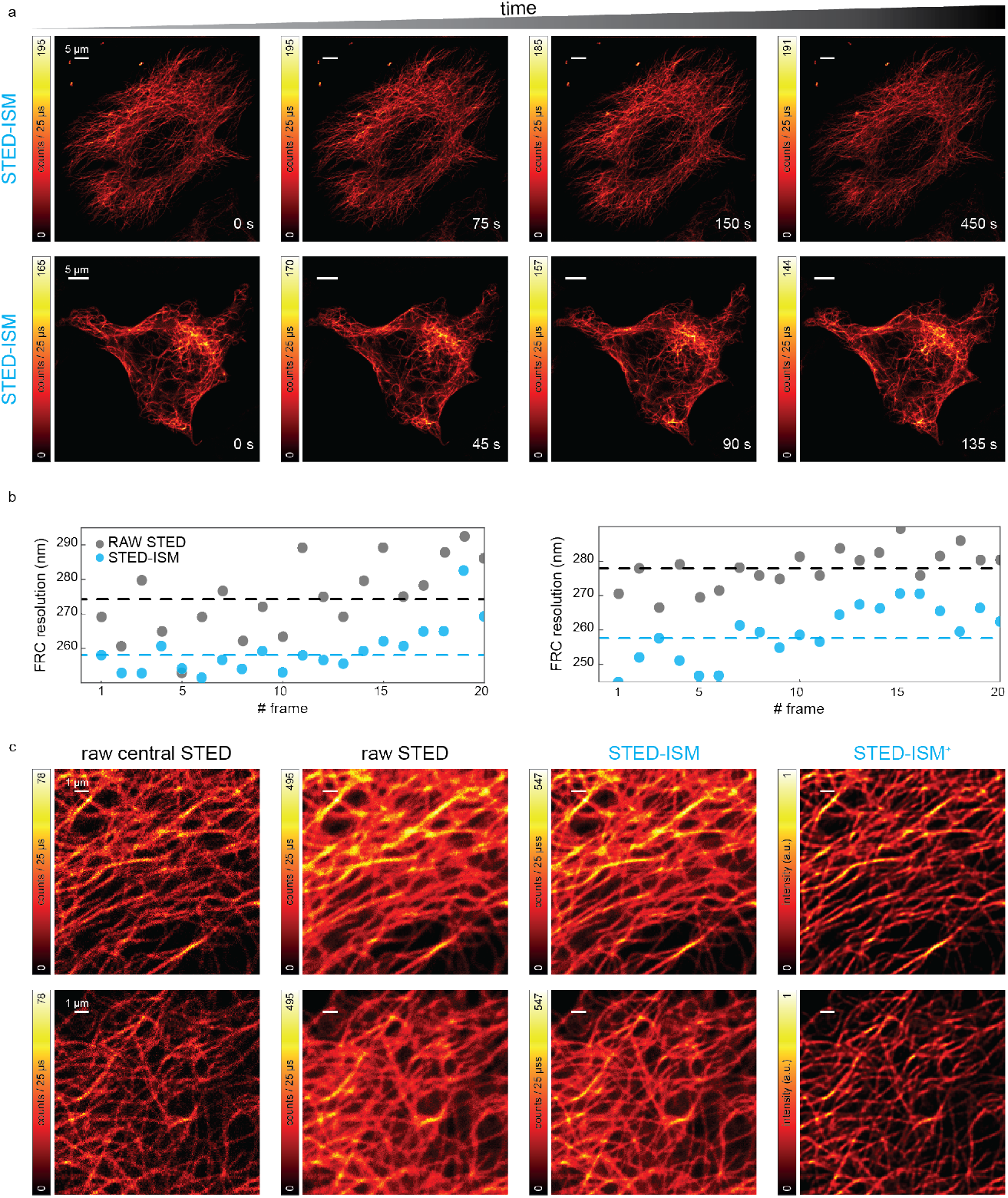
STED-ISM time series. **a**, Two extended time series (20 frames, 500 and 300 s) of Sir-Tubulin labeled Hela cells. Different frames are reported to show the negligible photo-bleaching. **b**, The FRC analysis confirms, for both time series, the superior performance of STED-ISM compared to raw STED. **c**, Details from the time series, showing the image recorded by the central element of the SPAD array (raw central STED), the result of the summing operation (raw STED), the result of the pixel reassignment operation (STED-ISM), and the deconvolution image (STED-ISM^+^). Pixel dwell time: 25 μs. Pixel size: 70 and 67 nm. Format of the full images: 1000 × 1000 and 750 × 750 pixels. Format of the details: 200 × 200 pixels. 635 nm excitation laser power: 8 μW; 775 nm STED laser power: 21 mW. Scale bars of **a**: 5 μm. Scale bars of **c**: 1 μm.

**Figure S6:**
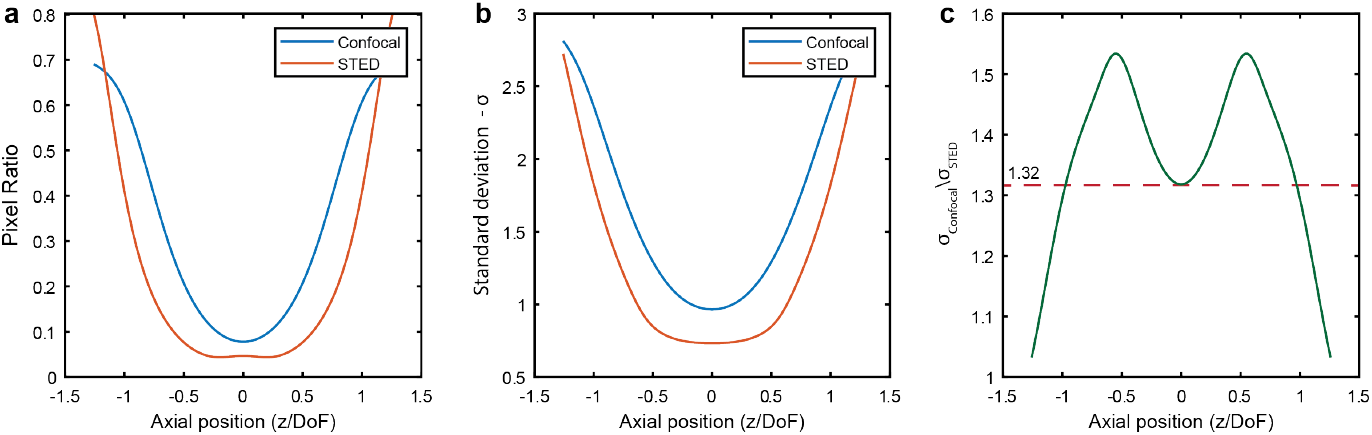
Distribution of light on the SPAD array. From the PSF simulations presented in Figure 4 we calculated a z-stack of fingerprints. In **a**, we show the ratio of the intensity averaged over the border elements to the intensity of the central element of the fingerprint. In **b**, we show the standard deviation resulting from a Gaussian fit to the fingerprints. In **c**, we show the ratio of the standard deviation of confocal fingerprint to the stadard deviation of the STED fingerprint. In the focal plane, the ratio is 1.32 to be compared with the theoretical expected value of 1.38. The axial units are normalized with respect to the depth-of-field (DoF = λ*n*/NA^2^).

**Figure S7:**
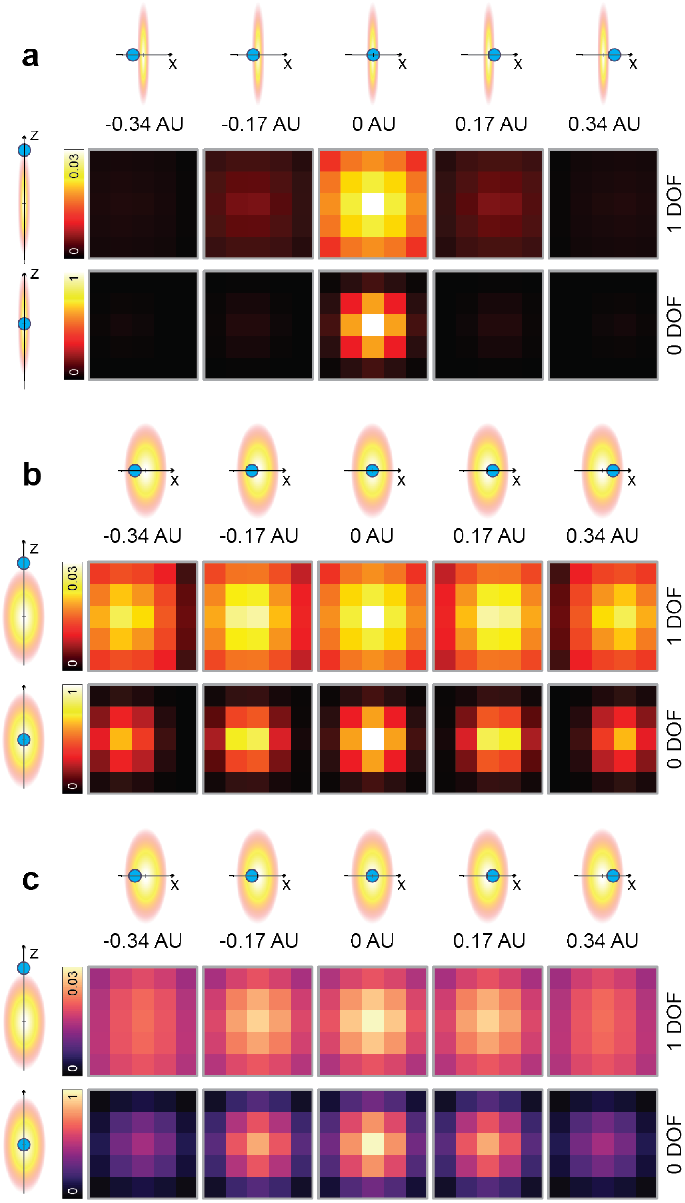
STED and confocal micro-images. Simulated micro-images generated by a single-emitter at different lateral and axial positions. In **a** we imposed ideal STED conditions (ς = 300). The micro-image corresponds to the detection PSF and contains a non-negligible amount of signal only if the emitter is centered in the *xy* plane. The lateral confinement occurs also out-of-focus. In **b**, we simulated the confocal case. The micro-images shift as the emitter moves laterally. In **c**, we show the micro-images from **b** after adaptive pixel reassignment. The new micro-images are now all centred and independent from the sample structure, except for a scale factor. In detail, they are proportional to the fingerprint of the corresponding axial plane. For visualization purposes, the simulatons are run with a detector size of one Airy unit.

**Figure S8:**
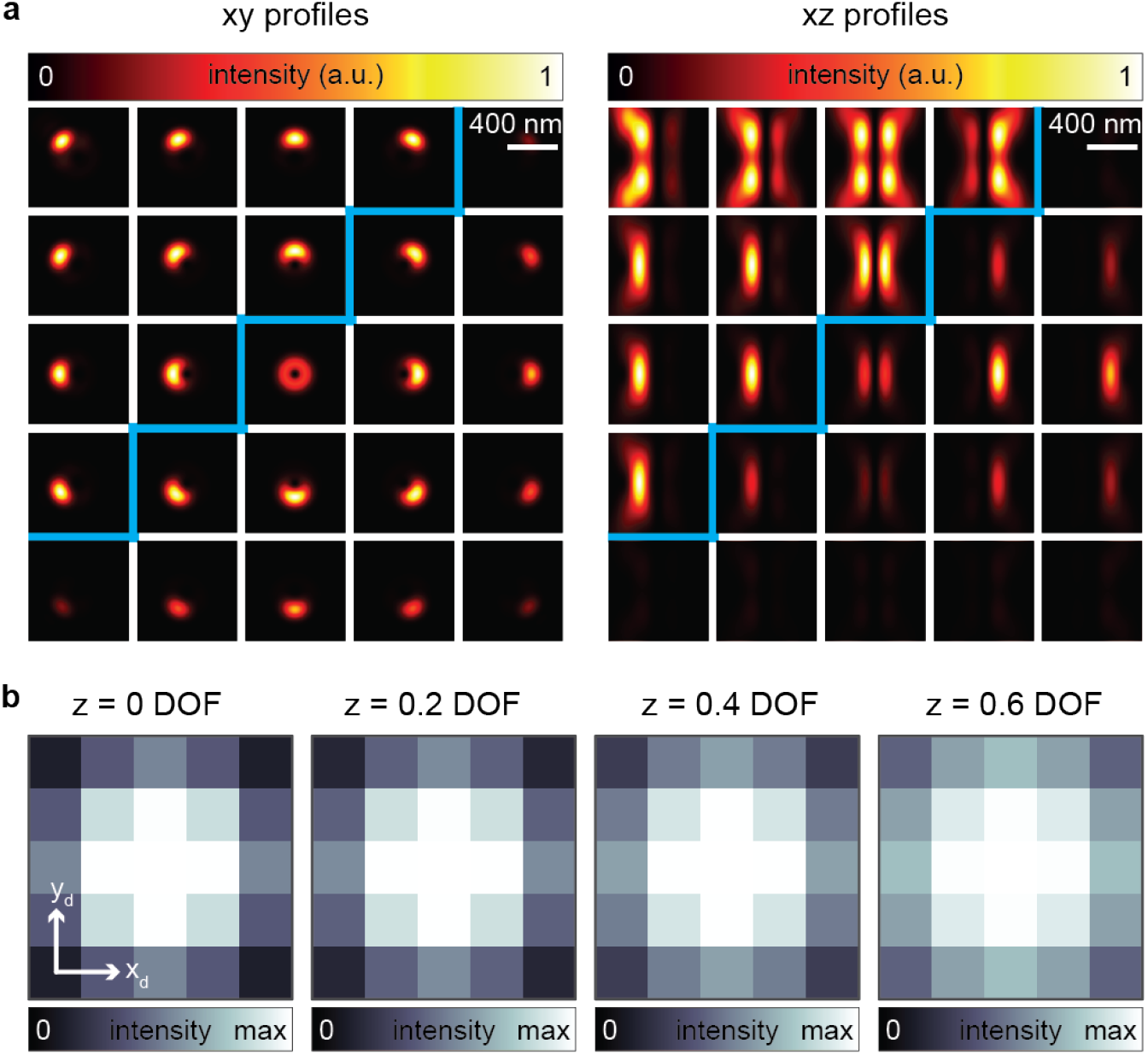
PSF in case of annular excitation. In **a**, we show the confocal PSFs obtained using an annular illumination. We calculated the PSFs for each detector element and for both the lateral and the axial plane. In **b**, we show the fingerprints calculated by summing all the scan points of the sections of the PSF at different axial positions. Notably, the intensity profile is flatter than in the case of Gaussian beam excitation.

**Figure S9:**
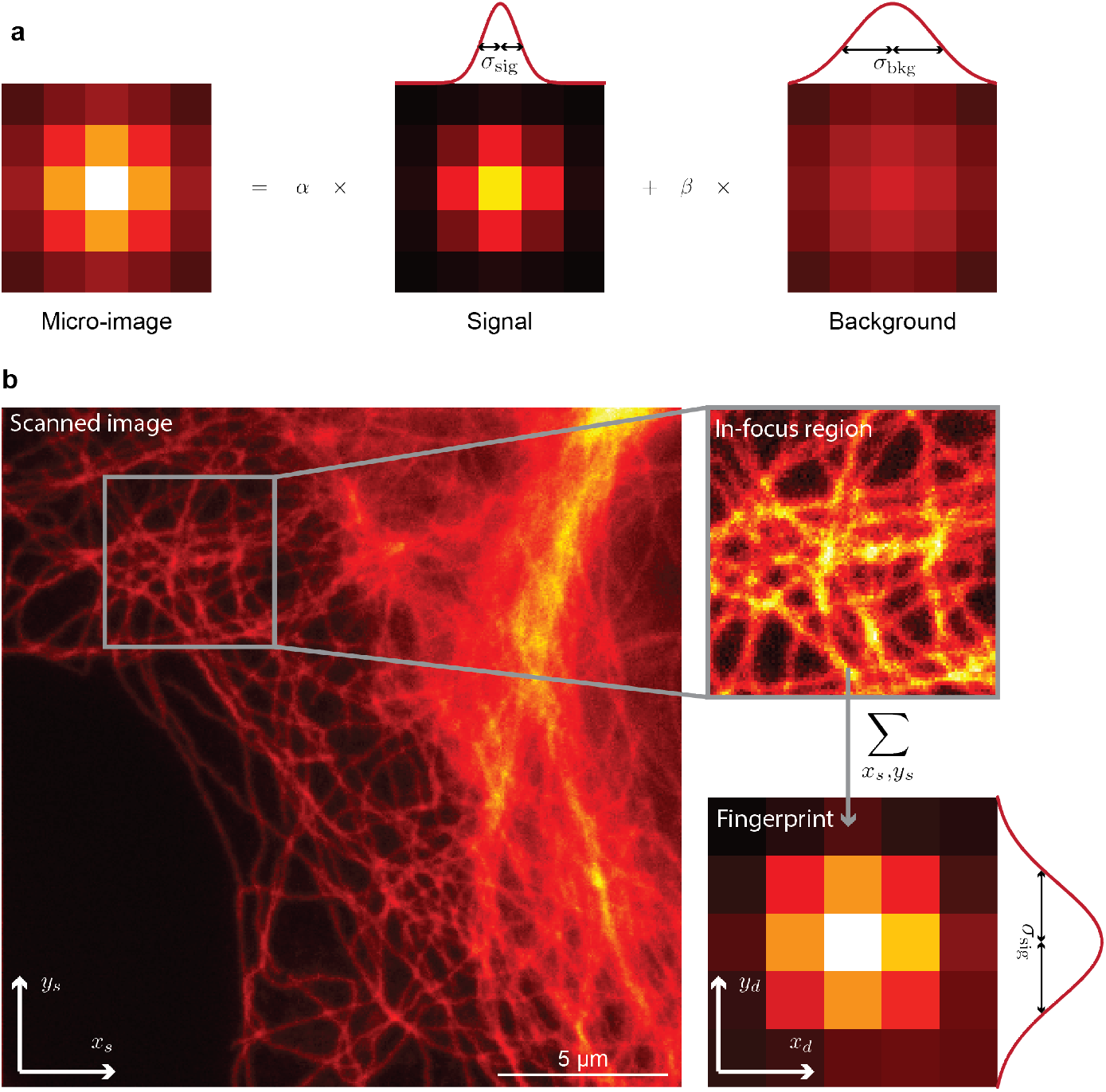
Working principle of the Focus-STED algorithm. In **a**, we show the core of the analysis: each micro-image is fitted to two Gaussian functions. The micro-image is normalized before the fitting and the weights respect the condition *α* + *β* = 1. The width of the in-focus Gaussian function is fixed either by using the theoretical value or by an additional measurement. In **b**, we show how to experimentally measure *σ*_sig_. We select a sub-image containing only in-focus emitters and calculate the corresponding fingerprint. The result is fitted to a single Gaussian function, whose standard deviation is used as *σ*_sig_ for the pixel-by-pixel fitting of the model presented in **a**.

**Figure S10:**
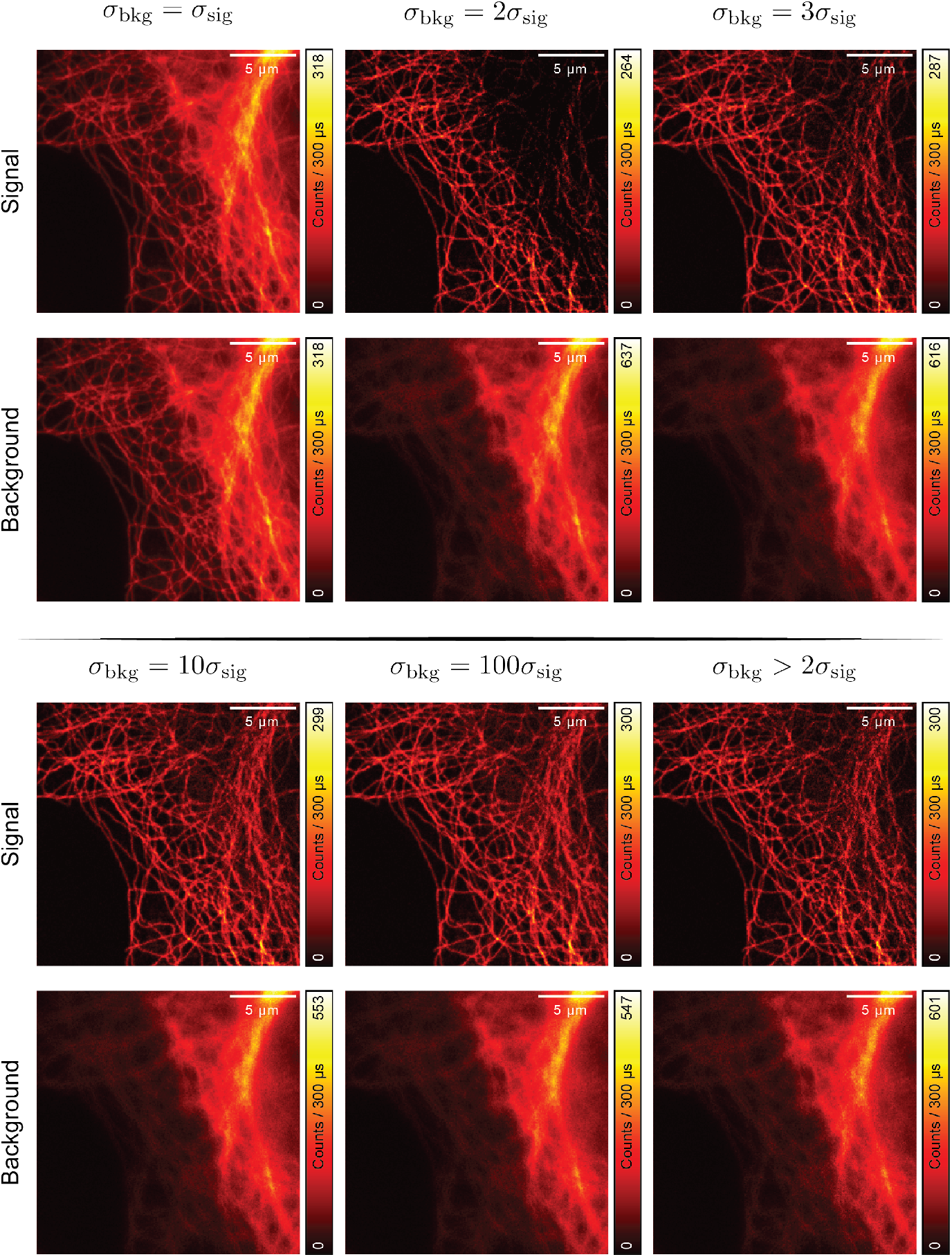
Result of the Focus-ISM algorithm. The first five reconstructions are obtained by fixing *σ*_bkg_. For *σ*_bkg_ = *σ*_sig_ the in-focus and out-of-focus photons follow exactly the same distribution and cannot be distinguished by the algorithm. Thus, the light is simply split into two identical images. For increasing *σ*_bkg_, the algorithm distinguishes the two contributions, but if *σ*_bkg_ is too small some misclassification may happen and too much signal is removed. For divergent *σ*_bkg_, the shape of the background approaches a constant offset. Notably, the result is still superior of that of the simple subtraction because of the physical constraint imposed by the fitting process. The last reconstruction is obtained letting *σ*_bkg_ be a fitting parameter, with the only physical constraint of being much larger than *σ*_sig_. With this last approach, *σ*_bkg_ is found adaptively pixel-by-pixel.

**Figure S11:**
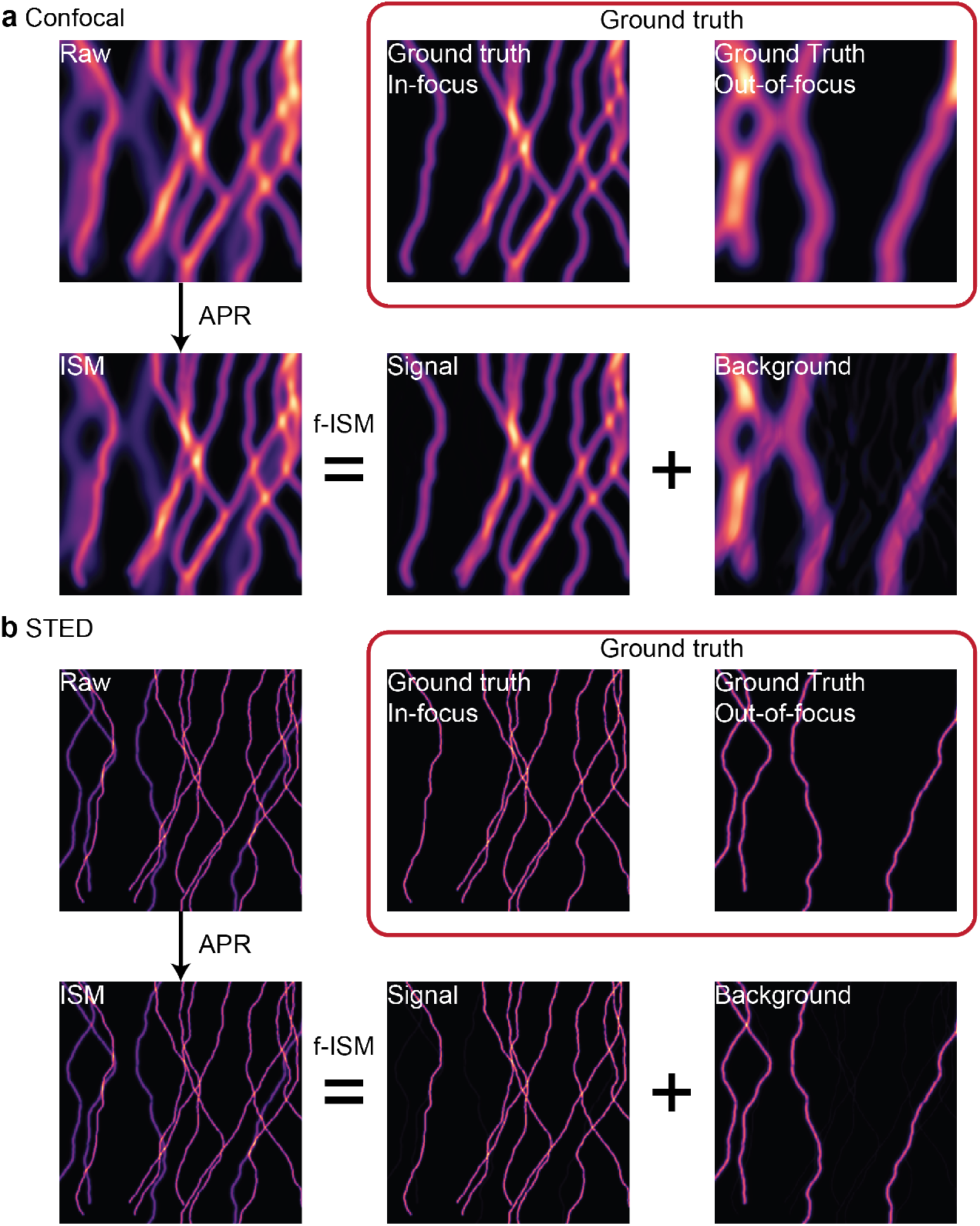
Result of Focus-ISM compared to the ground-truth. For each panel, we simulated tubulin filaments in-focus (*z* = 0) and out-of-focus (*z* = 1 DoF). We summed the two planes, generating an image affected by background *raw*). Then, we performed adaptive pixel reassignment, generating the ISM image. Lastly, we analyzed with the *f*^2^-ISM algorithm each micro-image of the reassigned dataset. The result is two images, one containing only the in-focus component *signal*) and one containing only the out-of-focus component *background*). On the bottom, we show the *ground-truth* for each component. Both in the confocal (**a**) and STED case (**b**) the result is very close to the corresponding ground-truth.

**Figure S12:**
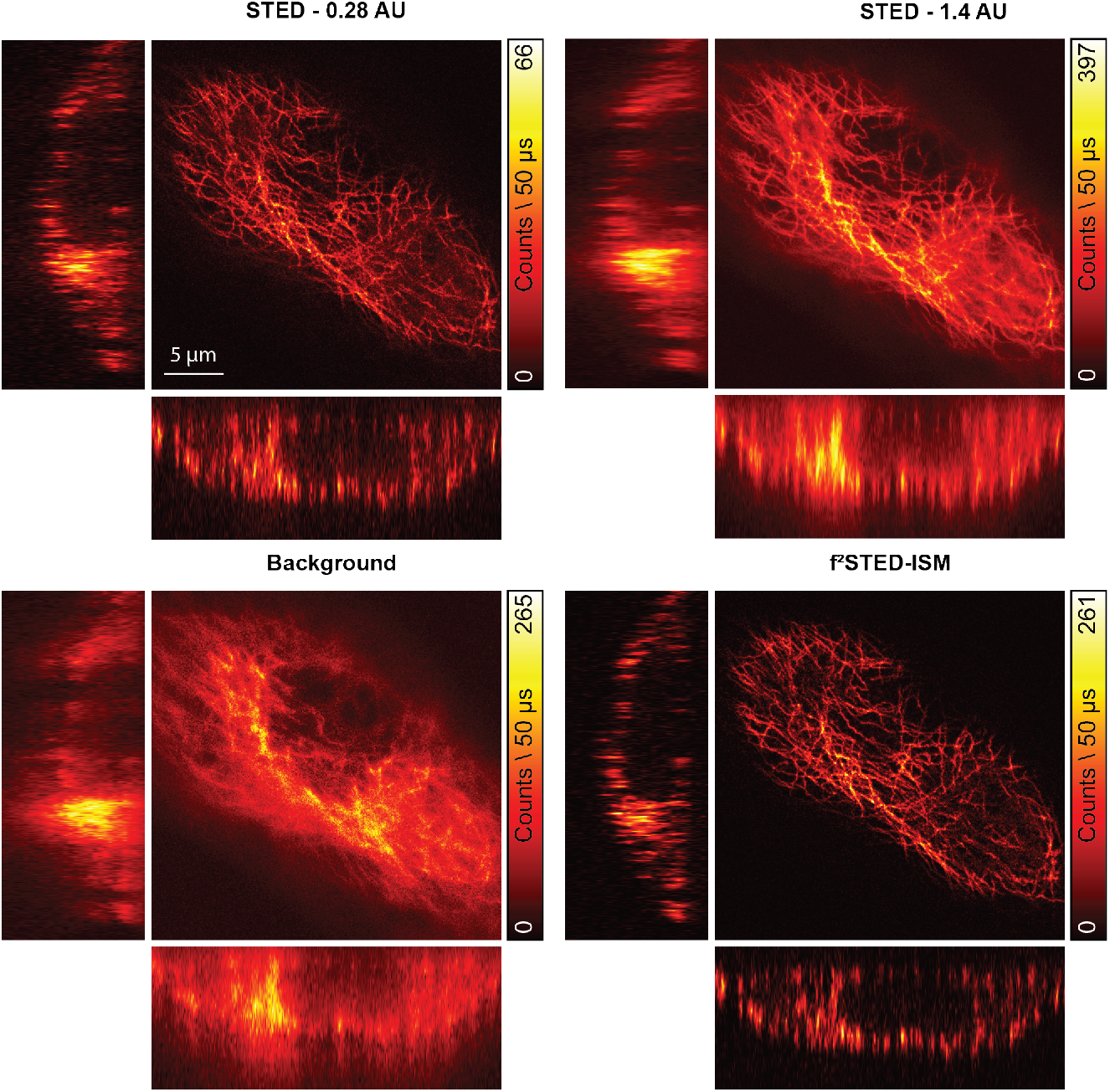
Focus-ISM applied to a z-stack of STED images. **Top left**, STED image acquired with a closed pinhole. The SBR is high, but the SNR is low. **Top right**, STED image acquired with an open pinhole. The SNR is high, but the SBR is low. **Bottom right**, result of the focus-ISM algorithm applied to the same dataset of the above images. Both SNR and SBR are high, and the similarity with the closed-pinhole image proves the absence of artifacts. **Bottom left**, image of the background removed. The *xz* and *yz* slices are taken from the middle of the volume and are linearly interpolated along the *z*-axis for visualization purposes. All images were acquired using an STED power of 125 mW measured before the objective lens.

**Figure S13:**
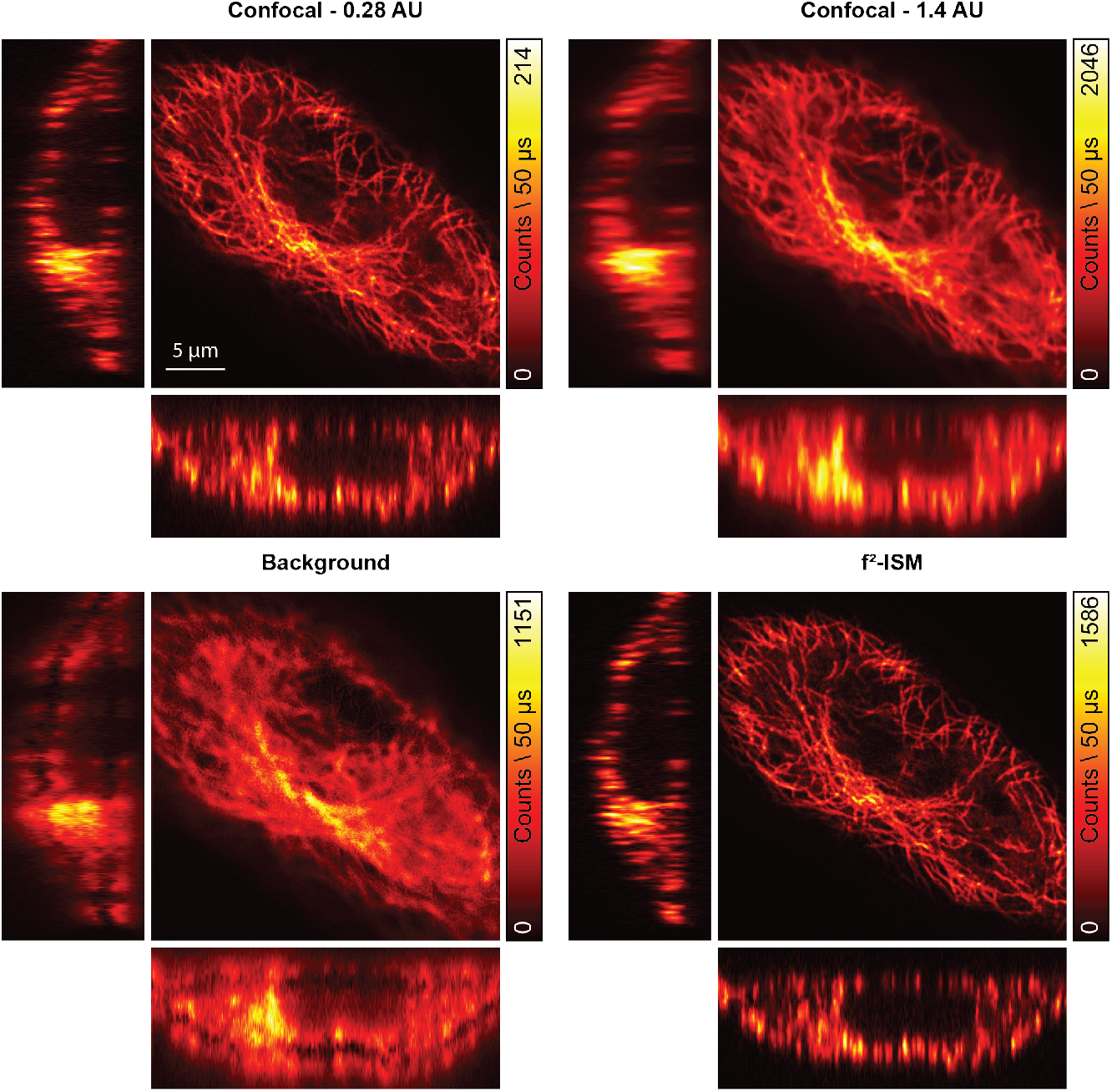
Focus-ISM applied to a z-stack of confocal images. **Top left**, confocal image acquired with a closed pinhole. The SBR is high, but the SNR is low. **Top right**, STED image acquired with an open pinhole. The SNR is high, but the SBR is low. **Bottom right**, result of the focus-ISM algorithm applied to the same dataset of the above images. Both the SNR and SBR are high, and the similarity with the closed-pinhole image proves the absence of artifacts. **Bottom left**, image of the background removed. The *xz* and *yz* slices are taken from the middle of the volume and are linearly interpolated along the *z*-axis for visualization purposes.

**Figure S14:**
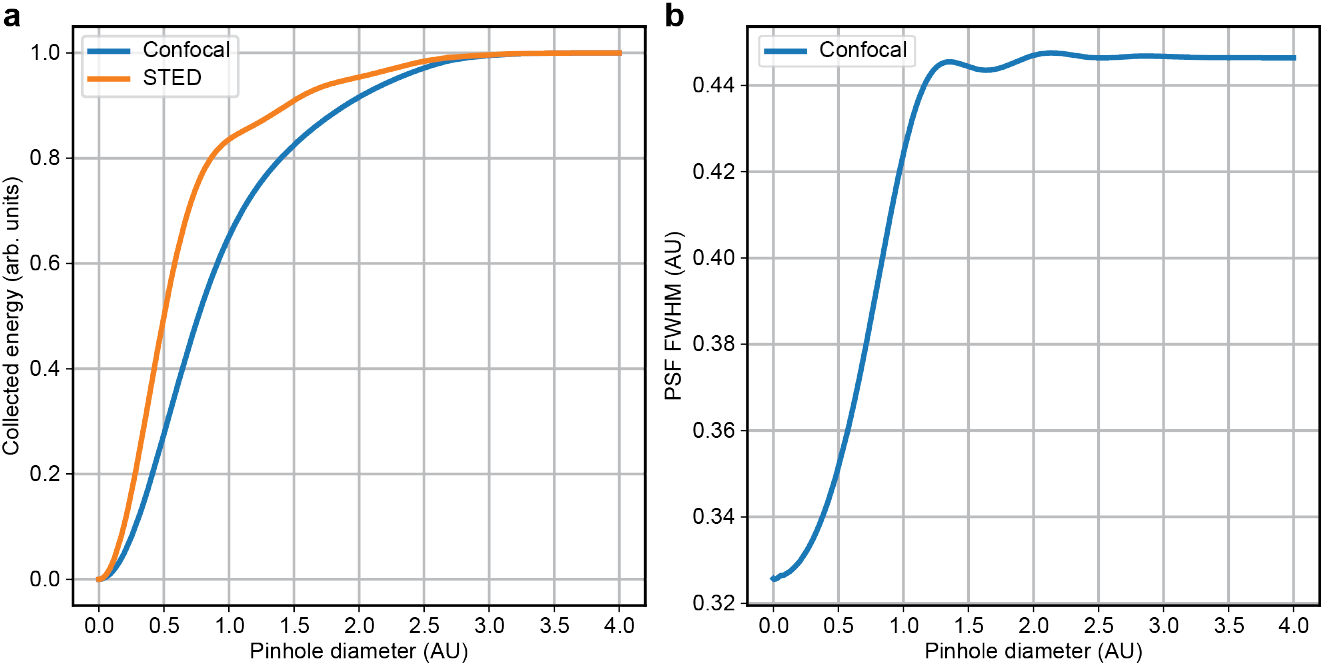
Collection efficiency at different pinhole sizes. In **a**, we show the results of the simulated collection efficiency from an in-focus sample at different pinhole size. In the STED case, the distribution of the light on the detector plane is given by the detection PSF. In the confocal case the light distribution is given by the convolution of the excitation PSF with the detection PSF, thus broader of a factor 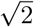 at the focal plane. Thus, the correct estimation of the correct pinhole size should consider the full fingerprint. The detection PSF is a an underestimation of intensity distribution on the detector, except than in the ideal STED case. In **b**, we show the full width at half maximum (FWHM) of the confocal PSF as a function of the diameter of the pinhole. The resolution and the collected signal are inversely proportional.

**Figure S15:**
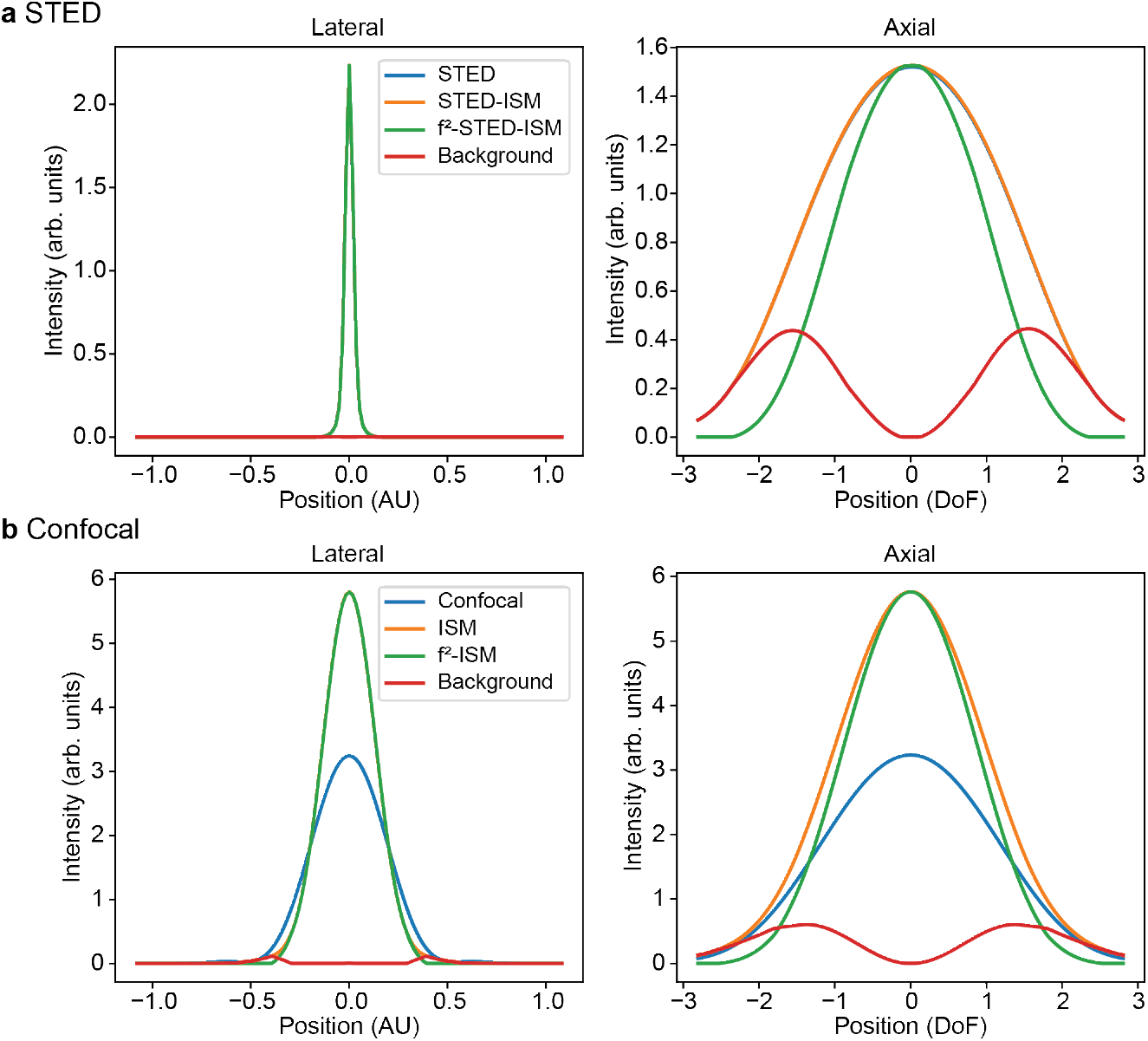
Intensity profiles of the reconstructed PSF. Intensity profiles of the PSFs shown in Fig. 4. Each line is extracted from the corresponding axis of symmetry of the PSF, namely *z* = 0 for the lateral profile and *x* = 0 for the lateral profile. In **a** we show the profiles of the STED PSFs (ς = 300), and in **b** we show the profiles of the confocal PSFs.

